# Microbiome-derived Short Chain Fatty Acids modulate microglial inflammatory responses in a sex- and metabolite-specific manner

**DOI:** 10.64898/2026.06.30.735602

**Authors:** Morgan Towriss, Vivien Dang, Jenna Goeres, Jatin Choudhary, Alison Aube, Juliana Montoya Sanchez, Clianta Anindya, Koyah Morgan-Banke, Jordan Hamden, Christopher Whidbey, Annie Ciernia

**Author notes:** Corresponding author Annie Vogel Ciernia, 2215 Wesbrook Mall, Vancouver, BC, V6T 1Z3. co-first authors (Morgan Towriss, Vivien Dang).

## Abstract

Microbes residing in the gastrointestinal tract exert immunomodulatory impacts on the brain through the gut-brain axis. Short-chain fatty acids (SCFAs) produced by bacterial fermentation of dietary fiber can enter the brain parenchyma and are implicated in microglia-mediated inflammation. While the gut microbiome is required to maintain microglial homeostasis, the mechanisms by which microbiota-derived metabolites affect microglia remains unknown. We examined the roles of SCFAs, specifically butyrate, propionate and acetate, on microglial function in response to SCFAs both *in vitro* using BV2 cells and *in vivo* in mice. We observed *in vivo* that SCFAs impact microglial transcriptional responses to LPS in a sex- and metabolite-specific manner with butyrate having the strongest effect. Enriched gene sets included signatures associated with LPS-responsive microglia, Arg1-positive microglia, microglial cell cycle related genes and genes affiliated with changes in microglial morphology. We observed a similar effect *in vitro,* where metabolite administration enhanced phagocytosis, blunted proliferation and nitric oxide production. We then evaluated global histone modification levels following metabolite treatment and detected an enhancement of H3K9ac, H3K27ac, and H3K4me3 both *in vivo* and in BV2 cells treated with butyrate. Finally, we showed that butyrate is a potent HDAC inhibitor possibly contributing to enhanced acetylation. Hence, our findings suggest that SCFAs impact microglial function in a metabolite- and sex-specific manner, and that butyrate blunts inflammation by regulating microglial histone acetylation. Our results provide a more in-depth understanding of gut microbiome-microglia crosstalk, opening the door for new microbiome- and microglia-targeted therapies.

**Highlights:** Xxxx

## 1. Introduction

Microglia are the resident macrophages of the central nervous system (CNS) and serve as the primary defense for the brain against injury and infection. In adulthood, microglia function to regulate synaptic pruning, cell death and homeostasis in the brain parenchyma (Hong et al., 2016; Nimmerjahn et al., 2005; Paolicelli et al., 2011; Tay et al., 2017; Thion and Garel, 2017). In response to pathological stimuli, microglia transition to a pro-inflammatory state and induce inflammation, which can be either protective or detrimental to the surrounding cells (Glass et al., 2010; Saijo and Glass, 2011). Microglial dysfunction or aberrant inflammatory programming has been linked to multiple neurological disease conditions, including schizophrenia, autism spectrum disorder (ASD), and Alzheimer’s disease (AD) (Colonna and Butovsky, 2017; Derecki et al., 2014; McCarthy and Wright, 2017; Volk, 2017). As such, regulation of microglial inflammation requires a complex combination of signals derived from their environment (Gosselin et al., 2017, 2014; Hickman et al., 2013; Lavin et al., 2014; Thion et al., 2018). The gut microbiome, acting through the gut–brain axis, provides peripheral environment signals that shape microglial development and function (Diaz Heijtz et al., 2011; Erny et al., 2015; Morais et al., 2021; Thion et al., 2018). Accordingly, the absence of the microbiome in germ-free (GF) mice or in dysbiosis contributes to microglial dysfunction and aggravation of disorders including AD or Parkinsons disease (PD) (Castillo-Ruiz et al., 2023, 2018; Hsiao et al., 2013; Liu et al., 2015; Sampson et al., 2016; Sullivan et al., 2025; Thion et al., 2018). Interestingly, the mechanisms linking microglial dysfunction to dysbiosis remain largely unclear. Bi-directional communication in gut-brain axis may include vagal nerve activity, cytokine signalling, immune cell trafficking, endocrine signalling and microbiota-derived metabolites capable of traversing or signalling through the blood brain barrier (BBB)(Braniste et al., 2014; Morais et al., 2021; Oldendorf, 1973). Recently, microbiota-derived metabolites have emerged as important modulators of microglial inflammation, yet the precise microbial sources, metabolites involved, and mechanisms of action remain poorly understood (Dalile et al., 2019; Den Besten et al., 2013).

- SCFA administration modulates microglial responses to immune challenge
- Microglia respond to SCFAs in a sex- and metabolite- specific manner
- Butyrate enhances microglial histone acetylation through HDAC inhibition

Short chain fatty acids (SCFAs) are one of the best studied classes of gut microbiota derived-metabolites that have been shown previously to traverse the BBB and directly impact CNS function (Dalile et al., 2019; Den Besten et al., 2013; Lu et al., 2016). Microbiota ferment dietary fiber in the colon, generating SCFAs which act both locally to maintain gut health and distally in peripheral tissues, including the CNS (Caetano and Castelucci, 2022; Caetano-Silva et al., 2023; Castillo-Ruiz et al., 2018). Among the diverse SCFAs produced by the commensal bacteria, butyrate, propionate and acetate have been studied most intensively due to their high abundance in the colon and their potential roles as key signalling molecules (Den Besten et al., 2013). Previous studies have demonstrated that intestinal dysbiosis, such as that induced by inflammation in murine models of inflammatory bowel disease (IBD), alters microglial morphology, density and gene expression (Caetano-Silva et al., 2023; Salvo et al., 2020; Sullivan et al., 2025). Notably, our previous work revealed that a reduction in SCFA producing microbes in an IBD model contributes to sex-dependent alterations in microglial activity (Sullivan et al., 2025). Further building on this literature, SCFA supplementation has been suggested as a therapeutic avenue for dysbiosis-related inflammation in the context of neurodegenerative diseases (Colombo et al., 2021; Erny et al., 2021; Fernando et al., 2020; Govindarajan et al., 2011; Seo et al., 2023). However, different SCFA treatments can have differential impacts on microglial activity and disease pathology, suggesting that individual SCFAs may act through different mechanisms to modulate microglial function.

How SCFAs regulate microglial function is still uncertain; evidence suggests they can either dampen or exacerbate microglia-mediated neuroinflammatory responses. Several different murine models of inflammation incorporating SCFA supplementation, suggest SCFAs exert anti-inflammatory effects on microglia (Caetano-Silva et al., 2023; Erny et al., 2021; Fernando et al., 2020). In addition, *in vitro* studies found that SCFAs reduce translocation of the pro-inflammatory NF-κB transcription factor into the nucleus, dampening expression of pro-inflammatory cytokines (Huuskonen et al., 2004; Wenzel et al., 2020). Conversely, multiple studies - particularly many *in vitro* investigations - suggest a pro-inflammatory effect of SCFA supplementation. For example, in the murine N9 microglial cell line, Huuskonen et al. observed that SCFAs potentiated lipopolysaccharide (LPS)-induced release of IL-6, which is canonically known as a pro-inflammatory cytokine (Huuskonen et al., 2004). Variability in findings may arise from context- and sex- specific differences in experimental models or distinct mechanisms by which SCFAs influence microglia (Hanamsagar et al., 2017; Markle et al., 2013; Sullivan et al., 2025; Thion et al., 2018). Additional investigation is required to clarify the mechanisms driving these divergent outcomes.

In addition to their impacts on cell signalling, SCFAs have been implicated in shaping the microglial epigenomic landscape by modulating histone acetylation (Caetano-Silva et al., 2023; Mann et al., 2024; Thion et al., 2018; Thomas and Denu, 2021). Histone acetylation is controlled by acetyltransferases (HATs) that add acetyl groups to histone tails, promoting transcription, and histone deacetylases (HDACs) that remove the acetyl groups, promoting transcriptional repression. Butyrate, in particular, has been reported to inhibit histone deacetylase 3 (HDAC3) in peripheral macrophages (Schulthess et al., 2019; Silva et al., 2018); HDAC3 inhibition has been shown to enhance microglial inflammatory cytokine expression and facilitate phagocytosis of debris and damaged cells (Meleady et al., 2023; Schulthess et al., 2019). A recent investigation by Thomas et al. revealed that in addition to blocking HDAC activity, butyrate and propionate were rapidly metabolized to the corresponding acyl-CoAs, activating the HAT p300 (Thomas and Denu, 2021). Consistently, SCFAs have been shown to modulate microglial gene expression and metabolic state by altering epigenetic marks including H3K4me3, H3K9ac and H3K27ac (Chen et al., 2024; Erny et al., 2021; Xie et al., 2026). Importantly, SCFA modulation of the microglial epigenome extends beyond simply blocking inflammatory responses and this complexity may contribute to the divergent outcomes observed across different studies.

Despite growing interest, conflicting evidence leaves the role of SCFAs in regulating microglial gene expression and function unresolved (Churchward et al., 2023; Colombo et al., 2021; De Witte et al., 2025; Fernando et al., 2020; Govindarajan et al., 2011; Schulthess et al., 2019; Wenzel et al., 2020). As such, this study employs both *in vivo* and *in vitro* models to provide a holistic view of SCFAs immunomodulatory influence on microglia. *In vivo,* we found that SCFA treatment blunted sickness responses to LPS injection, enhanced microglial histone acetylation, and blunted microglial inflammatory gene expression and morphology changes in a sex- and metabolite-specific manner. Further, different SCFAs had differential effects, indicating distinct roles for acetate, propionate and butyrate in microglial regulation with the largest impacts driven by butyrate. In BV2 immortalized microglial cells, SCFA administration reduced microglial proliferation, and enhanced phagocytosis while impairing NO release, with butyrate having the strongest effect. Our findings indicate that butyrate specifically acts as an HDAC inhibitor, suggesting a mechanism in which it prevents histone deacetylation, modulating gene expression to reduce inflammatory responses while enhancing microglial clearance of pathogens, ultimately facilitating resolution of inflammation. Together, these studies highlight the mechanistic effects of butyrate, propionate, and acetate on microglia, contributing to a more in-depth understanding of how the gut microbiota modulate microglial regulation.

## 2. Materials and Methods

### 2.1 In-vitro Methods

BV2 immortalized mouse microglia (Accegen #4125) were cultured in DMEM/F12 (ThermoFisher, Gibco #11320033), 10% HI-FBS (ThermoFisher, #12484028), 1X L-Glutamine (ThermoFisher, #25030081), and 1X Penicillin-Streptomycin (ThermoFisher, Gibco #15140148). All cells used for experiments were from passage 9-15. Before plating, cells were dislodged with 0.25% Trypsin- EDTA (ThermoFisher #25200072) and spun at 500g for 5 minutes, then plated in a 24-well plate with reduced serum media DMEM/F12, 2% HI-FBS, 1X L-Glutamine, and 1X Penicillin-Streptomycin.

#### 2.1.1 SCFAs and LPS treatments

Sodium butyrate (STEMCELL Technologies, #72242), sodium propionate (Sigma Aldrich, P5436), and sodium acetate (Sigma Aldrich, S5636) were dissolved and stored in sterile Phosphate-buffered Saline (PBS) (ThermoFisher, #14190144). BV2 cultures were pre-treated with 250 μM of each SCFA or with PBS as a vehicle control for 1 hour. Lipopolysaccharides (LPS) from *Escherichia coli* (Sigma Aldrich, #L5418) was diluted in H_2_O and added at a concentration of 10 ng/mL for 24 hours.

#### 2.1.2 Phagocytosis assay quantified by flow cytometry

BV2 cells were treated with SCFAs for 1 hour, LPS for 24 hours then for 1 hour with 500 ng/mL of pHrodo Red *E. coli* BioParticles (ThermoFisher Scientific, #P35361). Cells were trypsinized and moved to a 96-well round bottom plate (Corning, #3788) then fixed with 1% formaldehyde (Sigma Aldrich, F8775) for 20 minutes. FACS buffer was prepared from 1X Hank’s Buffered Salt Solution (HBSS) (ThermoFisher, #14185052), 1% Bovine Albumin Serum (BSA) (Tocris, #5217), 2 mM EDTA (ThermoFisher, AM9260G), and added to the cells for washing twice. The cells were then run on CytoFLEX Flow Cytometer, and data was analyzed by gating for cell size and granularity (FSC-A vs SSC-A), singlets (FSC-H vs FSC-width), and phycoerythrin (PE) (Y585) signal for detection of cells engulfing beads (Figure S8E). FlowJo was used to evaluate the percentage of bead positive cells gated on the no bead control in the PE channel.

#### 2.1.3 Global histone modification marker quantification by flow cytometry

BV2 cells were detached and moved to a 96-well round bottom plate upon completion of treatment as described above. Global histone modification level was assessed via flow cytometry as previously described (Towriss et al., 2023a)using the True-Nuclear Transcription Factor Buffer Set (Biolegend, #424401). Briefly, fixed cells were blocked with 10% Normal Donkey Serum (NDS) (Sigma Aldrich, D9663) and 1:100 TruStain FcX^TM^ PLUS (anti-mouse CD16/32) Antibody (Biolegend, 156604) for 30 minutes then stained with 250 ng/mL 4’,6-Diamidino-2-Phenylindole (DAPI) (Biolegend #422801) and primary antibodies for 30 minutes: 1:100 Acetyl-Histone H3 (Lys27) Rabbit mAb (Cell Signaling Technology, #8173), 1:200 Acetyl-Histone H3 (Lys9) Rabbit mAb (Cell Signaling Technology, #9649), or 1:400 Tri-methyl-histone H3 (Lys4) Rabbit mAb (Cell Signaling Technology, #9751). After two washes with 1X Permeabilization Buffer (PB), 1:200 Alexa Fluor 568 Donkey anti-Rabbit (Invitrogen, #A10042) was added as secondary antibody for 30 minutes. Cells were washed with 1X PB twice and resuspended in FACS buffer to be analyzed using CytoFLEX Flow Cytometer. FlowJo was used to gate the cells for cell size, granularity, singlets, and then for positive signal in both V450 channel for DAPI and mCherry (Y610) channel to detect antibody fluorescence. Median fluorescent intensity (MFI) of the positive population in Y610 channel was used as a measure for global histone modification level. MFI fold change was determined by comparing with vehicle conditions (MFI treated samples/ MFI vehicle samples). Gating strategy is provided in Figure S8C

#### 2.1.4 Griess reagent assay

BV2 cells were plated in a 24 well-plate with 300 μL of reduced serum media (2% FBS) made from DMEM/F12 without phenol red (ThermoFisher, #21041025). The experimental cells were plated for 24 hours prior to treatment as described above. The supernatant was then collected to quantify nitrite concentrations using the Griess Reagent Assay (ThermoFisher, #G7921). Sample absorbances were measured by a plate reader at 548 nm. Sample concentrations were then determined from a standard curve prepared from nitrite-containing standards. Lower limit of detection for this assay is 1.0uM.

#### 2.1.5 BrdU (Bromodeoxyuridine) cell proliferation assay

5-Bromo-2’-deoxyuridine (Toronto Research Chemicals, B683250) was dissolved in 1XPBS and sonicated for 10 minutes at 50°C. BV2 cells were treated with SCFAs and LPS as described and treated with 10μM BrdU for the last 3 hours of the LPS treatment. Cells were then fixed with 1% PFA and washed twice with 1X PBS 0.01% Tween 20 (ThermoFisher, BP337). Cells were treated with 1 M Hydrochloric acid (HCl) (ThermoFisher, A144-212) for 10 minutes to denature the DNA followed by a 30-minute neutralization with 0.1 M Sodium borate, made from Boric acid (Sigma Aldrich, B7901) and Sodium hydroxide (ThermoFisher, #BP359). Cells were blocked with 10% NDS, 5% BSA, and 22.52 mg/mL Glycine (Sigma Aldrich, G7126), and then stained with 50 ng/mL DAPI and 1:100 PE-Cyanine7 anti-BrdU Antibody (Biolegend, 364118) for 30 minutes. After two washes with 1X PBS, cells were resuspended in FACS buffer and run on CytoFLEX Flow Cytometer. FlowJo was used to gate the cells for cell size, granularity, singlets, and then for positive signal in PacBlue (V450) channel and PC7 (Y763) channel to detect Dapi+ cells with BrdU antibody fluorescence. Fold change in the BrdU positive population was determined by the percentage of DAPI and BrdU positive cells in treated samples divided by the same percentage in vehicle controls. Gating strategy is included in Figure S8D.

#### 2.1.6 Multiplex analysis of cytokines

BV2 cells were plated and treated with SCFAs and LPS as described above. Culture supernatants were collected after 24 hours for determination of released cytokines level. Secreted cytokine protein concentrations were measured using the Luminex xMAP technology for multiplexed quantification of 10 Mouse cytokines, chemokines and growth factors. The multiplexing analysis was performed using the Luminex™ 200 system (Luminex, Austin, TX, USA) by Eve Technologies Corp. (Calgary, Alberta). Ten markers were simultaneously measured in the samples using Eve Technologies’ Mouse Focused 10-Plex Discovery Assay® (MilliporeSigma, Burlington, Massachusetts, USA) according to the manufacturer’s protocol. The 10-plex consisted of GM-CSF, IFNγ, IL-1β, IL-2, IL-4, IL-6, IL-10, IL-12p70, MCP-1, and TNFα. Assay sensitivities of these markers range from 0.4 – 10.9 pg/mL for the 10-plex. Individual analyte sensitivity values are available in the MilliporeSigma MILLIPLEX® MAP protocol.

#### 2.1.7 XTT viability assay

In a 96-well plate, 10 000 BV2 cells were plated in 100 μL of reduced serum media (2% FBS) made from DMEM/F12 without phenol red (ThermoFisher, #21041025) for 24 hours before treatment with SCFAs and LPS as described above. 70 μL of XTT working solution provided in the CyQUANT^TM^ XTT Cell Viability Assay (ThermoFisher, #X12223) was added to each well. After 4 hours, the absorbance at 450 nm and 660 nm was read using a plate reader. Dead control samples were treated with 0.5% Triton-X100^TM^ (FisherScientific, #BP151-500) for 30 minutes while blank controls were prepared with only XTT reagent and no cells. The cell viability fold change was calculated as the difference between absorbance measurements (450 nm – 660 nm), corrected for absorbance of the blank and dead controls, then divided by the PBS-treated samples.

#### 2.1.8 HDAC-Glo I/II screening assay

10,000 BV2 cells were seeded into a 96-well plate in 50 μL of reduced serum media (2% FBS) made from DMEM/F12 without phenol red (ThermoFisher, #21042015) and incubated overnight. In parallel, two-fold serial dilutions of each SCFA and Trichostatin A (positive control) were prepared in a final volume of 75 μL of HDAC-Glo^TM^ Buffer (Promega, G6420). Negative controls included without inhibition (No Metabolite) and without HDAC (No Cell) samples. Reduced culture medium was then replaced with 50 μL of serum-free medium followed by a 30-minute incubation with 50 μL of prepared serial dilutions. 100 μL of HDAC-Glo^TM^ Reagent (Promega, G6420) was added to each well, and the plate was incubated at room temperature for 30 minutes. Luminescence was measured by a plate luminometer (RLU). The percent inhibition was calculated as (1 − 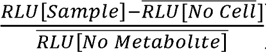) × 100. For analysis, values below 0% or above 100% inhibition were constrained to 0% and 100%, respectively, to improve IC50 curve-fitting stability. IC50 values were calculated in GraphPad Prism using four-parameter nonlinear regression.

### 2.2 In vivo Methods

Specific pathogen free (SPF) experimentally naïve 8–10-week-old male and female C57BL6J mice (strain #000664, Jackson Laboratories) were bred in house and housed across litters in groups of 2-4 under a standard 12:12 light-dark cycle with sex and treatment-matched cage mates. All experiments were performed during the mouse’s regular light cycle. Animals had *ad libitum* access to food (Teklad 2918) and sterile reverse osmosis chlorinated water enriched with salt solutions as described below. Animals were given standard amounts of beta chip bedding and nestlet, crinkle paper and red hut enrichment. All housing conditions and testing procedures were in accordance with Canadian Council on Animal Care guidelines, and all protocols were approved by the Animal Care Committee of the University of British Columbia (A23-0086).

#### 2.2.1 In vivo experimental design

Mice of each sex were assigned to four groups by cage and administered sterile pH-balanced isotonic solutions of either: (i) Saline (67.5 mM NaCl (ThermoFisher, P358 212)); (ii) Butyrate (40 mM sodium butyrate (Sigma Aldrich, S303410)), (iii) Propionate (25 mM sodium propionate (Sigma Aldrich, P5436)), or (iv) Acetate (67.5 mM sodium acetate (Sigma Aldrich, S5636))(Colombo et al., 2021) (Table 1). For solution preparation, sterile water from our animal facility was used to dissolve the salts; the solutions were then filtered (0.22μm) prior to being added into the facility standard water bottles under sterile conditions. Water solutions were given *ad libitum* and replaced every 3-4 days for 4 weeks. The animals were regularly monitored for water consumption, body conditioning, and behavioral changes. After 4 weeks, the animals underwent behavioral testing prior to further manipulation.

**Table 1.** Final concentrations of SCFA water solutions.

| Condition | SCFA concentration (mM) | NaCl concentration (mM) |
| --- | --- | --- |
| NaCl vehicle | 0 | 67.5 |
| Butyrate | 40 | 27.5 |
| Propionate | 25 | 42.5 |
| Acetate | 67.5 | 0 |

Within cage, the animals were assigned at random to receive either 1 mg/kg LPS (Sigma Aldrich, #L5418) or physiological saline (VWR #470302-026) resulting in a total of 8 groups per sex (Saline PBS, Saline LPS, Butyrate PBS, Butyrate LPS, Propionate PBS, Propionate LPS, Acetate PBS, Acetate LPS). Animals were injected intraperitoneally between 09:00 and 11:00 on the day prior to tissue collection. Animals were scored for sickness behavior three hours after the injection.

#### 2.2.2 Behavioral testing: Open Field test

All behavior testing was performed under white light (lux=190) between the hours of 11:00 and 15:00 after 3-4 weeks of treatment to assess behavioral effects of the SCFA treatment. Open field test was performed in an open-topped grey plexiglass arena light chamber (65 cm x 65 cm x 40 cm; *lwh*). Mice were placed in the center of the area one at a time and allowed to freely explore for 10 minutes. ANY-maze software (Stoelting) was used to track the total distance travelled and the amount of time spent in the center zone within the arena. The zone boundary was defined at 10cm from the apparatus wall. The percentage of time spent in the center of the field was used as a metric for animal anxiety, and a two-way ANOVA was performed to test for effects of SCFAs and Sex. Tukey-adjusted post hoc comparisons were performed between treatments.

#### 2.2.3 Water consumption and body weight change analysis

Body weights and water levels were measured every 3–4 days, coinciding with the refreshing of the water solutions. Water consumption was calculated as the change in bottle weight (in mL) since the previous measurement. Outliers within each sex and diet condition were identified and removed using a 1.5× interquartile range (IQR) filter. The cleaned data were analyzed using a linear mixed-effects model with Diet, Sex and Day as fixed effects and Cage as a random intercept to account for repeated measures. An additional fixed effect of Cohort was included to control for batch variation. To assess whether SCFA treatments influenced water consumption over time, slope comparisons across diets within each sex were performed using estimated marginal trends (emtrends() from the emmeans package). Tukey-adjusted post hoc comparisons were conducted to identify significant differences in water consumption slopes between Diet groups.

To analyze body weight, outliers within each sex, diet and LPS injection group were identified and removed using a 1.5× interquartile range (IQR) filter. The cleaned dataset was analyzed using a linear mixed-effects model implemented in the lmerTest and nlme packages in R. The model included Diet, Sex, LPS Injection, and Day as fixed effects, with Animal as a random effect to account for repeated measures. An additional Cohort effect was included to account for batch variability. To assess whether SCFA treatments influenced weight trajectories over time, slope comparisons across diets within each sex and LPS injection condition were performed using estimated marginal trends (emtrends() from the emmeans package). Tukey-adjusted post hoc comparisons were conducted to identify significant differences in weight change slopes between Diet groups.

#### 2.2.4 Sickness behavior scoring

Changes in sickness behavior were assessed 3 hours after the injection. Sickness behavior was scored following a 4-point grading scale (0 to 3 points total) as previously described ((Kim et al., 2024b). Lethargy signified by decreased mobility and curved body posture, ptosis signified by facial grimace and drooping eyelids, and piloerection signified by ruffled, greasy fur and poor grooming were each assessed individually for either 0, 0.25, 0.5, 0.75 or 1 point. A cumulative score of 0 would suggest that no symptoms of sickness behavior were present whereas a cumulative score of 3 points indicates that all symptoms of sickness behavior were maximally present. Sickness was scored 3 hours after the injection timepoint, and researchers were blinded to animal condition at the time of scoring. Change in sickness score was assessed by fitting a non-parametric Aligned Ranks Transformation (ART) model to measure scores as a factor of SCFA and LPS treatments using the ARTool R package. The model was assessed using ART ANOVA and post hoc tests between treatment groups were corrected for multiple comparisons using the Holm method.

#### 2.2.5 Microglia isolation

Mice were anesthetized with isoflurane and transcardially perfused with 1X PBS with inhibitors cocktail, including 5 μg/mL Actinomycin D (NEB, 15021S), 10 μM Triptolide (NEB, 97539), and 27.1 μg/mL Anisomycin (NEB, 2222S)(Marsh et al., 2022; Towriss et al., 2023a). Brains were extracted and minced into small (<1 mm) pieces in 1.5mL tubes using a scissor. Minced samples were then digested with 800 μL digestion buffer per brain (100 μg/mL DNase I (STEMCELL Technologies, 07900), 20 units/sample activated Papain solution (Worthington Biochemical, LS003124), and the inhibitor cocktail). Brains were digested at 37°C for 30 minutes at 660rpm shaking. Digested brain solution was transferred into a 15 mL glass dounce homogenizer on ice with 5 mL FACS buffer and dounced until homogenous. Brain homogenate was transferred to new 15 mL tubes, where 2.125 mL of 100% Percoll solution (Millipore Sigma, GE17-0891-02) was added and FACS buffer was used to top up until 8.5 mL of total volume. 4 mL of 37% Percoll and 2 mL 70% Percoll solutions were then underlaid respectively. Layered gradients were centrifuged at 500xg for 20 minutes at 4°C at the centrifuge’s minimum brake setting. Myelin and supernatant layers were discarded, leaving 2 mL of the interlayer containing microglia between 37% and 70% gradients. The interlayer was topped up with 10 mL FACS for washing and centrifuged at 500xg for 10 minutes at 4°C with full brake on. Finally, supernatant was removed, and cell pellets were resuspended in 200 μL remaining FACS buffer before being transferred to 1.5 mL tubes for staining.

##### 2.2.6.1 RNA-sequencing library preparation and sequencing analysis

Total RNA was extracted from isolated microglia using the RNeasy Plus Microkit (Qiagen, 74134) 24 h after systemic LPS challenge in male and female mice maintained for 4 weeks on saline, acetate, propionate, or butyrate-supplemented diets. Samples were analyzed for RNA Integrity Scores greater than 9 and 2-10ng total RNA. 18-20 PCR cycles were used to generate Poly-A sequencing libraries using Illumina Stranded mRNA prep (Illumina). Libraries were pooled and sequenced on an Illumina NextSeq2000 with Paired End 59bp × 59bp reads to an average read depth of 29 million reads. Sequencing data was demultiplexed using Illumina’s BCL Convert. De-multiplexed read sequences were then aligned to the Mus Musculus (mm10) reference sequence and transcripts counted per gene using DRAGEN RNA app on Basespace Sequence Hub (https://support-docs.illumina.com/SW/DRAGEN_v41/Content/SW/DRAGEN/TPipelineIntro_fDG.htm).

##### 2.2.6.2 Weighted Gene Co-expression Network Analysis (WGCNA)

Gene co-expression network analysis was performed using the **WGCNA** R package to identify transcriptional modules associated with diet, LPS challenge, and sex in isolated microglia. RNA-seq count data were imported into R genes with low expression were removed using filterByExpr, retaining genes expressed in at least 50% of samples within experimental conditions. Counts were converted to log2 counts per million (log-CPM) using a prior count of 1. To reduce noise, genes with variance below the 25th percentile were excluded. Batch effects attributable to cohort were removed using limma’s removeBatchEffect function while preserving biological variables (diet, injection, sex, and their interactions). The resulting expression matrix contained 16,403 genes across 83 samples. Data quality was assessed using goodSamplesGenes, and hierarchical clustering of samples revealed no outliers requiring removal (Figure S2A).

A signed co-expression network was constructed using biweight midcorrelation (bicor) to reduce sensitivity to outliers. Soft-thresholding power was evaluated across a range of values (1–30) using pickSoftThreshold (Figure S1B). Network construction and module detection were performed using blockwiseModules with signed topology, minimum module size of 25 genes, reassignment threshold of 0, and a maximum proportion of outliers set to 0.05. Modules were identified using hierarchical clustering of the topological overlap matrix (Figure S2C). Module eigengenes (MEs) were calculated as the first principal component of each module (Figure S2D). Associations between module eigengenes and experimental traits (Diet, LPS treatment, Diet × LPS interaction, and Sex) were assessed using Pearson correlations, with statistical significance evaluated using Student’s t-tests. Resulting p-values were adjusted for multiple comparisons using the false discovery rate (FDR) across the full correlation matrix.

Gene ontology enrichment analyses for individual modules were performed using clusterProfiler against a universe of all genes included in the WGCNA input. Enrichment was assessed separately for Biological Process, Molecular Function, and Cellular Component ontologies using Benjamini–Hochberg correction. Enrichments for microglia specific gene lists was performed using MGEnrichment with FDR correction (https://ciernialab.shinyapps.io/MGEnrichmentApp/) (Jao and Ciernia, 2021). Enrichment results were visualized using dot plots scaled by odds ratio and –log10(FDR).

##### 2.2.6.3 Differential gene expression analysis

Raw gene-level counts from two sequencing batches were combined as raw count matrices, imported into R and processed using edgeR and limma. A single sample (SCFA2.11) was excluded due to an apparent dosing inconsistency. Lowly expressed genes were filtered using filterByExpr, retaining genes expressed in at least 50% of samples within experimental conditions. Library sizes were normalized using trimmed mean of M-values (TMM). Log2 counts per million (CPM) were calculated, and data distributions were assessed via density plots and library size visualizations (Figure S3A and B). Gene-wise linear modeling was performed using the limma-voom framework. A design matrix was constructed modeling each sex × diet × injection condition explicitly while including experimental cohort as a covariate to control for batch effects. The voom function was used to estimate the mean–variance relationship and generate observation-level precision weights. Model fit diagnostics were assessed via voom mean–variance plots (Figure S3C). Sample relationships were visualized using multidimensional scaling (MDS) plots based on normalized expression data, both before and after batch correction using removeBatchEffect. Batch-corrected expression values retained biological condition effects while minimizing cohort-driven variation (Figure S3D).

Contrasts were defined to test: (i) LPS versus PBS responses within each diet and sex, (ii) diet effects within PBS or LPS conditions, and (iii) sex-collapsed effects computed as the average of male and female contrasts. Linear models were fit using lmFit, contrasts applied with contrasts.fit, and empirical Bayes moderation performed using eBayes. Differentially expressed genes (DEGs) were identified using Benjamini–Hochberg false discovery rate (FDR) correction, with genes considered significant at adjusted *p* < 0.05. To compare sex-specific transcriptional responses, DEGs identified in male, female, and sex-collapsed contrasts were compared using UpSet plots generated with UpSetR. Overlaps among up- and down-regulated gene sets were assessed within each diet condition. Euler diagrams were generated to visualize shared and sex-specific DEG sets for LPS-responsive genes using the eulerr package.

##### 2.2.6.4 Rank–rank hypergeometric overlap (RRHO2)

Ranked overlap between differential gene expression signatures was assessed using Rank-Rank Hypergeometric Overlap (RRHO2) in R. For each LPS versus VEH comparison, genes were ranked by signed significance, calculated as −log10(adjusted *p*-value) multiplied by the direction of change. RRHO2 analyses were performed separately by sex to compare SCFA treatments (butyrate, acetate, or propionate) for LPS versus VEH gene lists. RRHO2 objects were generated using default parameters and overlap significance was visualized as heatmaps depicting regions of concordant and discordant regulation across ranked gene lists.

##### 2.2.6.5 Enrichment analysis (GSEA)

Gene set enrichment analysis (GSEA) was performed using ranked log2 Fold Change expression results derived from RNA-seq data. Microglia-specific gene sets were obtained from a curated mouse microglial gene list database from MGEnrichment (Jao and Ciernia, 2021), filtered to retain genes expressed in the dataset and gene sets containing ≥10 genes. GSEA was conducted in R using the *fgsea* package with default parameters (minimum gene set size = 10, maximum = 500). Enrichment results were generated for LPS vs PBS contrasts and Diet effects within LPS or PBS, stratified by Sex. Normalized enrichment scores (NES) and FDR–adjusted *p*-values were calculated for each gene set comparison. Significant enrichments (adjusted *p* < 0.05) were visualized using dot plots, where dot size represents enrichment magnitude and color reflects statistical significance.

#### 2.2.7 In vivo flow cytometry and analysis

Extracellular staining for cell sorting: This procedure involves staining of live cells and thus is performed on ice. Cell pellets from microglia isolation were blocked with 1:100 TruStain FcX^TM^ PLUS (anti-mouse CD16/32) diluted in FACS buffer for 10 minutes. Cells were then stained for 30 minutes with 1:100 Life Dead Zombie Aqua dye (Biolegend, 423101), 1:100 PE anti-P2RY12 (Biolegend, 392103), 1:200 APC anti-mouse CD45 (Biolegend, 103112), and 1:200 FITC anti-mouse/human CD11b (Biolegend,101205). The live dead (L/D) control was created by snap freezing 50% of cells in this sample. Cells were then washed twice with FACS, then resuspended in 500 μL FACS buffer, and strained through a 70 μm reversible filter (STEMCELL Technologies, 27216) prior to sorting using CytoFLEX SRT sorter. Cells were gated based on size, granularity, singlet, and signals in KRO-V525 channel (LD stain), FITC-B525 channel (CD11b), Y585 channel (P2RY12) and APC-R660 channel (CD45). Microglia were collected from live cell population that was CD11b+, CD45 low and P2RY12+ (Fig S8A).

Intracellular staining for histone marks: Resuspended cells after being stained for extracellular markers (CD45, CD11b, P2RY12) were transferred to a 96-well round bottom plate. Global histone modification levels were assessed via flow cytometry using the True-Nuclear Transcription Factor Buffer Set as described above. Cells were blocked with 5% Normal Donkey Serum (NDS) for 10 minutes. Cells were then incubated with 1:50 Pacific Blue^TM^ Conjugate Acetyl-Histone H3 (Lys9) Rabbit mAb (Cell Signaling Technology, #11857) and 1:50 PE Conjugate Acetyl-Histone H3 (Lys27) Rabbit mAb (Cell Signaling Technology, #15562) for 30 minutes. After two washes with 2% NDS diluted in 1X PB provided in the kit, cells were resuspended in FACS buffer and run on CytoFLEX Flow Cytometer. FlowJo was used to gate the cells for cell size, granularity, singlets. Microglia were determined from CD11b +, CD45 (low) in the FITC-B525 and APC-R660 channels respectively. From identified microglia population, cells were gated for H3K27ac (PE-Y585) and H3K9ac (PacBlue -V450). Median Fluorescence Intensity (MFI) of the positive population in the relevant channel was used as a measure for global histone modification level. MFI fold change was determined by comparing with vehicle conditions (MFI of treated samples/ MFI vehicle samples) (Fig S8B).

#### 2.2.8 Immunofluorescence staining, imaging and morphology analysis

Mice received repeated intraperitoneal injections of either LPS or vehicle (PBS), with or without SCFA treatment, and brains were collected 24 hours after the final injection. All animals were cardiac perfused with PBS prior to brain collection. Brains were fixed in 4% paraformaldehyde for 48hrs at 4C, transferred to 30% sucrose and stored at 4C until frozen for sectioning on a cryostat at 30um. Tissue sections underwent antigen retrieval (Sections were incubated in 10mM sodium citrate and 0.05% Triton X-100 at 80C for 30 minutes, then for 10 minutes at room temperature) and then were permeabilized (1xPBS□and□0.5% Triton (Fisher BioReagents BP151-500) for 5 minutes), and blocked (1xPBS,□0.03% Triton,□and□1% Bovine Serum Albumin (BSA; Bio-techne Tocris 5217) for 1 h). Sections were incubated overnight at 4°C in primary antibody solution (guinea pig anti IBA1, Synaptic Systems 234 308, concentration) containing 2% Normal Donkey Serum (NDS; Jackson Immunoresearch Laboratories Inc. 017-000-121),□1xPBS,□0.03% Triton. After primary antibody incubation, sections were washed 3 times for 5 minutes each with 1xPBS□and□0.03% Triton before incubating for 2 h in secondary solution containing 2% NDS,□DAPI (1:1000; Biolegend 422801), and□secondary antibody (Donkey anti-Guinea Pig Alexa Fluor 647; Jackson ImmunoResearch # 706-605-148 1:500). Sections were washed 3 times for 5 minutes each with 1xPBS and□0.03% Triton and 1 time for 5 minutes with 1xPBS before being transferred into 1xPBS for storage before mounting. Sections were mounted onto microscope slides (Premium Superfrost Plus Microscrope Slides, VWR CA48311-703) and air dried before being coverslipped with mounting media (EverBrite Fluorescence Antifade Mounting Media, Biotium 23001). Mounted sections were imaged on a ZEISS Axioscan 6 microscope slide scanner at 20□×□magnification with a step-size of 1um using the z-stack acquisition parameters within the imaging software (ZEISS ZEN 3.7). Extended Depth of Focus (EDF) images were generated in ZEN software using maximum intensity projection, which compiles the brightest pixels across a z-stack to produce a 2D image that preserves key 3D structural information. The hippocampus subfields CA1, CA3 and Dentate Gyrus (DG), and prefrontal cortex (PFC) were selected as regions of interest (ROI) using the Allen Brain Atlas (mouse.brain-map.org) using the ImageJ macro FASTMAP (Terstege et al., 2022). ROIs were then used as input for analysis using the ImageJ plugin MicrogliaMorphology and R package MicrogliaMorphologyR toolsets (Kim et al., 2024a). The parameters used for MicrogliaMorphology included: auto local thresholding using the NiBlack and radius 100 with an area filter of 310.69–879.57 μm^2^. Morphology feature values were log-transformed and filtered for outliers using a 3× IQR rule. Principal component analysis (PCA) was performed on the 27 standardized morphology features to reduce dimensionality. The first three PCs were used as input for k-means clustering. Optimal cluster number was determined using within-cluster sum of squares and silhouette methods, with four clusters selected. Clusters were annotated based on morphology as ramified, ameboid, rod-like, or hypertrophic. Cluster distributions were analyzed separately for each region as percentages of microglia per cluster per mouse. A linear mixed-effects model was used to assess the effects of Cluster, Diet, LPS Injection, Sex, and their interactions, with MouseID as a random effect. Post hoc comparisons were corrected using Sidak adjustment.

### 2.3 General statistical analysis

For all experiments except weight change analysis, sickness behavior, and open field test, two-way ANOVA was used to fit a full effect model (LPS effect, SCFA effect, and their interaction) followed by Tukey’s post hoc analysis to compare individual treatments. All measures passed normality and homoscedasticity tests performed in GraphPad Prism 10.2.1. Outliers were eliminated using ROUT method (Q=1%).

## 3. Results

### 3.1 *In vivo* SCFA supplementation alters LPS-induced microglia in a sex dependent manner

To investigate the impact of SCFAs on microglia under physiological conditions, we supplemented adult C57BL6 mice with SCFA through drinking water (“Diet”) (Fig 1A) *ad libitum* for 4 weeks. Water consumption and mouse weight were monitored throughout the 4 weeks, and we found no significant impact of treatment on weight or water consumption (Fig S1A-B, Table S1). Additionally, we assessed if SCFA supplementation impacted mouse locomotion or anxiety using an Open Field Test and found no effects (Fig S1C, Table S1) (La-Vu et al., 2020; Seibenhener and Wooten, 2015). Together these data suggest that the diet supplementation was well tolerated and did not cause gross changes to mouse behavior. After four weeks of supplementation, the mice were injected with 1mg/kg bacterial lipopolysaccharide (LPS) or sterile saline (VEH). After 3 hours, animal sickness behavior was scored based on piloerection (0-1), ptosis (0-1) and lethargy (0-1) to yield a total sickness score (0-3). There was a main effect of both LPS (p<0.005) and Diet (p<0.005) and a significant Diet x Treatment x Sex interaction (FigS1D, Table S1). Posthoc comparisons revealed that LPS significantly increased sickness regardless of Diet, but butyrate significant blunted LPS-induced sickness behavior in both sexes, and propionate significantly blunted LPS-induced sickness in males. Together, these results suggest that SCFA supplementation is well tolerated and butyrate blunts sickness responses to LPS-induced inflammation.

**Figure 1:**
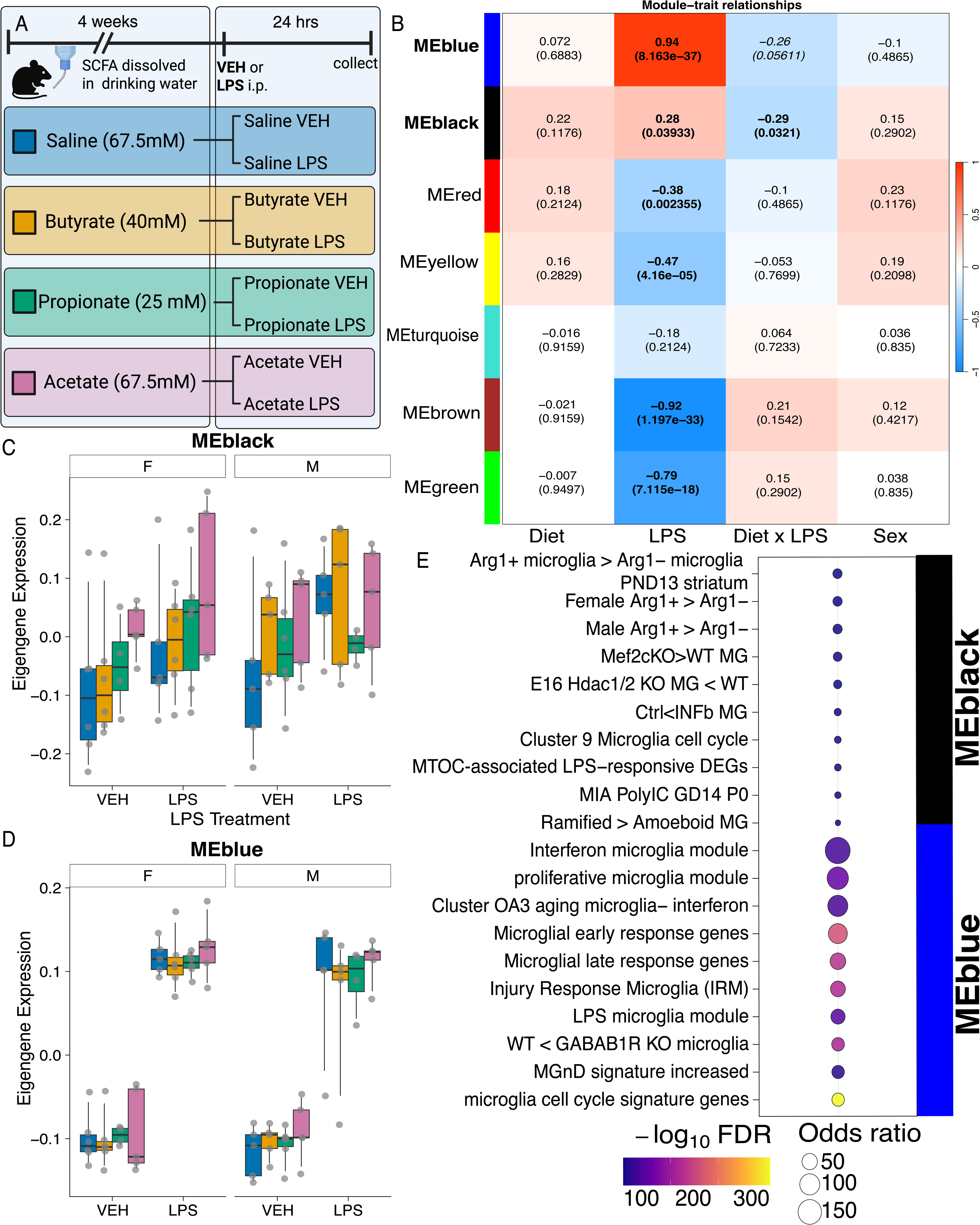
SCFA supplementation modulates gene cluster changes in response to LPS administration. (A) Outline of SCFA and LPS treatment paradigm in specific pathogen free C57Bl6/J mice. Mice were administered either Sodium Chloride (67.5mM), Sodium Butyrate (40mM), Sodium Propionate (25mM) or Sodium Acetate (67.5mM) in drinking water for 4 weeks. After 4 weeks, the animals were injected intraperitoneally with bacterial lipopolysaccharide (LPS) at 1 mg/kg or sterile saline (0.9%) (VEH) and collected 24 hrs after. Figure was made using biorender. (B) WGCNA Network. (C) Eigengene boxplot for the Black module and (D) Blue module. (E) Dot plot showing GSEA results for Blue and Black modules. Dot size reflects the Odds Ratio and color represents –log10 FDR adjusted p-values.

#### 3.1.1 SCFA supplementation suppresses microglial inflammatory programming in a sex- and metabolite-dependent manner

To examine the impacts of SCFA on LPS-induced microglial gene expression, microglia were isolated using flow cytometry (CD45+/Cdllb+ cells) from combined cortex and hippocampus, and RNA was collected for RNA-sequencing. To examine coordinated gene expression programs underlying SCFA- and LPS-dependent transcriptional changes in microglia, we performed weighted gene co-expression network analysis (WGCNA) on batch-corrected, variance-filtered log2-CPM expression data from all samples (Fig S2, Table S2). Correlation of module eigengenes with experimental variables revealed several modules significantly associated with LPS treatment and the interaction of LPS and SCFA supplementation as after FDR correction (Fig 1B). No modules arose as significantly associated with SCFA supplementation alone. Among the modules, the *black module* exhibited a positive association with LPS and the strongest negative association with the Interaction, indicating sensitivity to inflammatory challenge that was negatively modulated by SCFA supplementation. Similarly, the *blue module* had the strongest positive association with LPS and a negative association with the Interaction that neared significance (FDR p=0.056). Examination of *black module* eigengene showed modulation of the VEH and LPS conditions by Diet in both sexes, while *blue module* showed modulation of the LPS response that was more pronounced in the males (Fig 1C-D).

Gene ontology and curated microglial enrichment analyses of *black module* genes revealed significant enrichment of pathways related to inflammatory activation, RNA processing, cytoskeletal organization, and immune signalling functions (Table S2). We further evaluated the *black module* genes using MGEnrichment, a custom database of published microglial gene lists and human neurologic disorders (Jao and Ciernia, 2021). Enriched gene lists within the MGEnrichment package included signatures associated with LPS-responsive microglia, Arg1-positive microglia, microglial cell cycle related genes and genes affiliated with changes in microglial morphology (Fig 1E, Table S2). Similarly, MGEnrichment analysis of *blue module* genes revealed gene signatures associated with the LPS response, interferon response and proliferative microglia (Fig1E, Table S2). Together, these results indicate that SCFA supplementation may modulate microglial inflammatory signalling through the complex regulation of a coordinated transcriptional program that integrates inflammation with cellular stress mechanisms.

To further parse the role of the individual SCFAs in regulating microglial inflammatory responses we performed differential gene expression analysis for each of the SCFA conditions. Differential gene expression analysis revealed no significantly differentially expressed genes (FDR < 0.05) between Diet supplementation conditions in either the LPS or VEH conditions (e.g. VEH Saline vs VEH Butyrate; LPS Saline vs LPS Propionate) (Table S3). However, there were significant effects of Diet on the response to LPS administration (e.g. VEH vs LPS-Saline vs VEH vs LPS-Butyrate) resulting in genes both increased and decreased after LPS administration unique to each Diet condition (Table S3, Fig S4A-H). Comparing gene lists between males and females for these contrasts showed that there were more differentially expressed genes arising from each Diet in females compared to males, especially for butyrate (Fig S4A-H, Table S3). Subsequently, comparing the contrasts between Diet conditions, revealed both a set of conserved genes that are regulated regardless of Diet and a set of genes regulated uniquely under each Diet condition supporting metabolite specific effects (Fig S4I-L). We then compared gene list regulation using Rank–rank hypergeometric overlap (RRHO2) analysis (Fig 2A-F) for LPS versus VEH -log(p-values) ranked gene lists within each Diet. SCFAs altered LPS-induced transcriptional responses in microglia in both sexes, as evidenced by prominent off-diagonal RRHO signals indicating shifts in gene regulation patterns between diet conditions. Butyrate produced the strongest and most coherent overlap pattern, particularly in females (Fig 2A,D), while acetate showed intermediate effects (Fig 2B,E) and propionate showed weaker and more heterogeneous impacts (Fig 2C,F). Together, these findings suggest that individual SCFAs differentially reshape microglial transcriptional responses to inflammatory challenge in a metabolite-dependent manner, with evidence for sex-specific effects.

**Figure 2:**
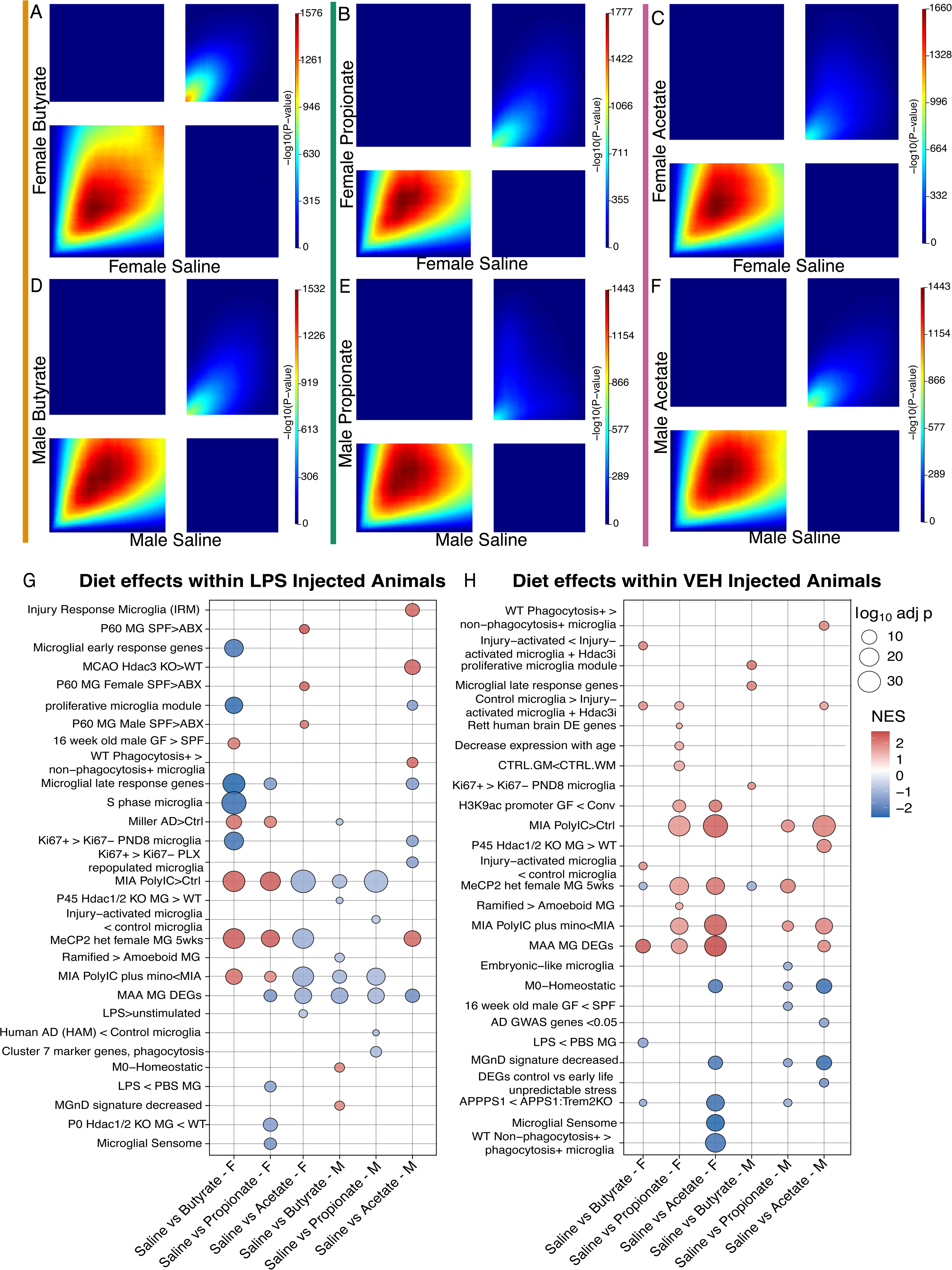
SCFAs induce sex- and metabolite-specific changes to microglia at baseline and under inflammation. (A-F) Rank–rank hypergeometric overlap (RRHO2) heatmaps showing genome-wide similarity in transcriptional responses to LPS between SCFA-treated and saline-treated microglia, stratified by sex and SCFA. Heatmaps depict overlap between -log(p-value) gene rankings from LPS versus VEH for each SCFA treated versus saline comparison in females (A–C) and males (D–F). Axes represent genes ranked by -log(p-values), with genes higher in SCFA conditions toward the upper/left extremes and genes higher in saline conditions toward the lower/right extremes. Color intensity reflects the statistical significance of overlap (–log10 hypergeometric p value). (G) Dot plot showing GSEA results for dietary SCFA supplementation under LPS challenge (LPS□+□SCFA versus LPS□+□saline), shown separately for females (F) and males (M). Pathways with higher expression in LPS□+□SCFA relative to LPS□+□saline have a positive normalized enrichment score (NES) (red), whereas pathways higher expression in LPS□+□saline have a negative NES (blue). Dot size corresponds to –log10 FDR adjusted p-values. (H) Dot plot showing GSEA results for SCFA supplementation under basal conditions (VEH□+□SCFA versus VEH□+□saline), shown for females (F) and males (M). Pathways with higher expression in VEH +□SCFA relative to VEH +□saline have a positive NES (red), whereas pathways higher expression in VEH +□saline have a negative NES (blue). Dot size reflects –log10 FDR adjusted p-values and color represents NES.

To further evaluate these more subtle shifts in gene expression by Diet, we performed Gene Set Enrichment Analysis (GSEA) using gene sets from MGEnrichment. Within the LPS condition, SCFAs modulated microglial transcriptional programs in a sex- and metabolite-specific manner relative to LPS□treated□saline animals (Fig 2G, Table S4). Butyrate supplementation showed the strongest suppression of inflammatory and disease-associated pathways, with several gene sets displaying higher expression in LPS□treated□saline than in LPS□treated□butyrate samples (negative NES, blue), including Injury-activated microglia, proliferative microglia, microglial late response genes, and disease-linked signatures such as human Alzheimer’s Disease microglia and microglial phagocytosis-positive states. Interestingly, these pathways were enriched strongly for female butyrate supplemented mice, but not for male, suggesting sex-specific programming. In contrast, pathways enriched in LPS□mice treated with□SCFA (positive NES, red), particularly within acetate treated males, included gene sets related to metabolic fitness and stress adaptation including pro-phagocytic, oxidative phosphorylation and Arg1-negative microglia. Propionate exhibited intermediate effects, with fewer pathways showing strong enrichment in either direction. In the absence of immune challenge (VEH treatment), SCFA supplementation reinforced homeostatic microglial transcriptional programs resulting in enriched pathways supporting morphology changes and phagocytic activity in microglia (Fig 2H, Table S4). The effects were generally weaker than those observed following LPS administration, consistent with the findings of the WGCNA analysis showing interaction effects in LPS responsive genes. Consistent with these findings, GSEA comparing LPS to VEH treatments directly within each Diet revealed that LPS increased inflammatory and disease-associated transcriptional programs (Fig S4M, Table S4), indicating that the impact of SCFAs is modulatory, not repressive of LPS induced inflammation. Together, these results indicate that under inflammatory challenge SCFAs do not uniformly impact microglial states but instead modulate inflammatory programming arising from blunted disease signatures (most evident for butyrate in females) and enhancing metabolic and adaptive responses (most evident for acetate in males).

#### 3.1.2 SCFA supplementation alters microglial morphology responses to LPS

As the transcriptomic analysis revealed enrichment in pathways associated with microglial morphology changes (*ramified vs amoeboid cells*), we hypothesized that SCFA supplementation *in vivo* would alter shifts in microglial morphology in response to LPS administration. Using the same model as described in Fig1A, tissue was collected 24 hrs after LPS administration and morphology analysis was performed as previously described (Kim et al., 2024a). Four morphology clusters were identified from the 58,241 microglia analyzed across two different brain regions (prefrontal cortex (PFC) and hippocampal (HPC) - subregions CA1, CA3 and DG) from both sexes and all treatments (Fig 3A, Fig S5, Table S5).

**Figure 3.**
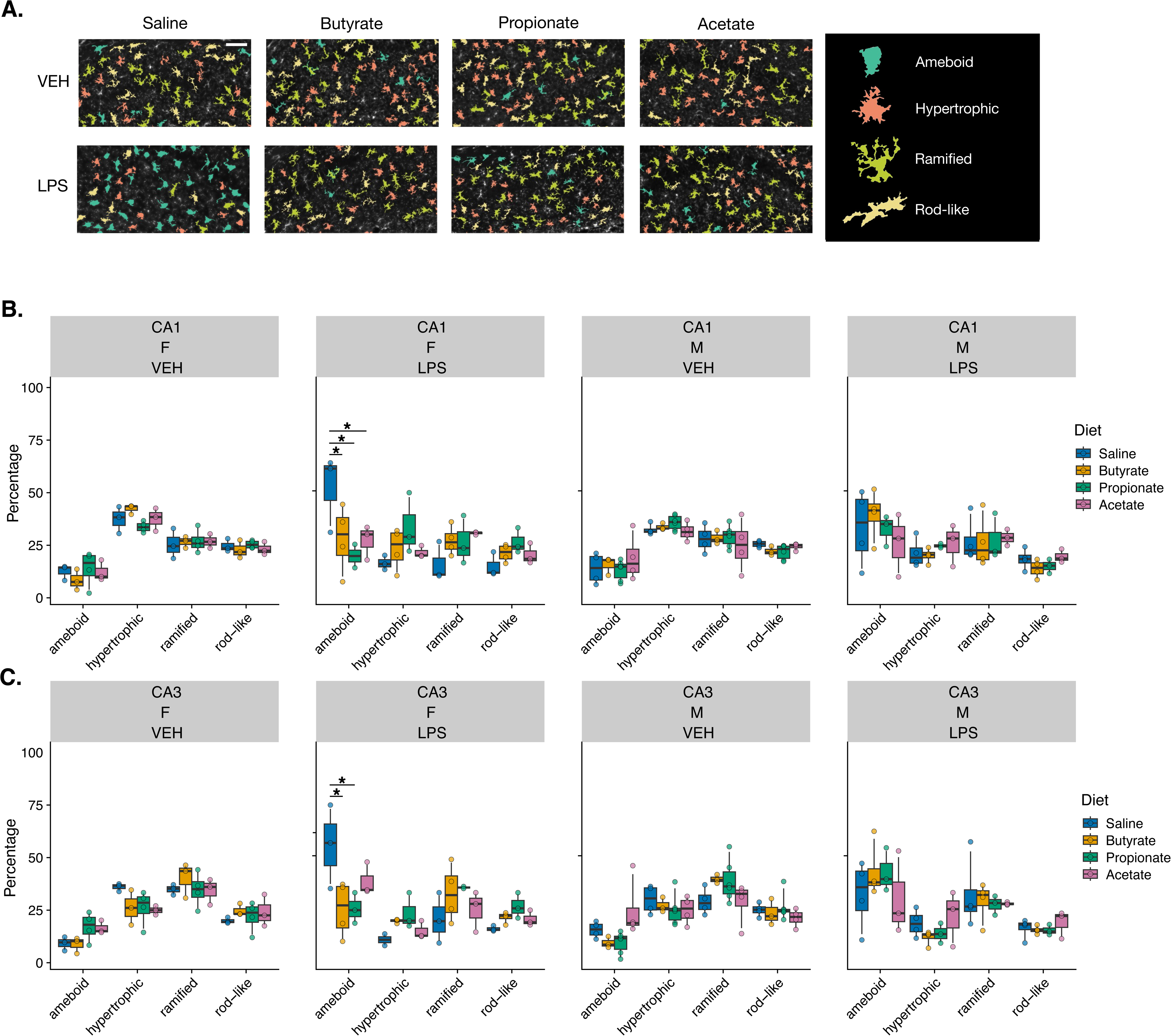
SCFAs blunt microglial morphology shifts in response to LPS in females. (A) Representative classifications of microglia into the 4 identified clusters. Cluster identification is overlaid on IBA1 stained images of CA1 microglia from each treatment group for females. (B) Female mice show a significant increase in ameboid microglia in response to LPS that is blunted by butyrate, propionate and acetate in CA1. Male mice show an LPS induced increase in ameboid microglia that is not blunted by SCFA treatment. (C) Female mice show a significant increase in ameboid microglia in CA3 in response to LPS that is blunted significantly by butyrate and propionate. Male mice show an increase in ameboid microglia in response to LPS that is not blunted by SCFA treatment. n=3-4 mice/sex/diet/treatment; 160-3635 microglia per region per mouse, average of 728 microglia/mouse/region. * p<0.05 Sidak corrected post-hoc comparisons.

In CA1 there was a significant interaction between microglial Cluster, Diet and Sex and a significant interaction between Cluster, Diet and LPS injection (Table S5). In accordance with previous studies, the saline treated mice showed a significant increase in proportion of ameboid microglia and decrease in hypertrophic microglia after LPS administration (Kim et al., 2024b). The LPS-induced increase in ameboid microglia was significantly blunted in the female mice treated with butyrate, propionate and acetate, but not the male mice (Fig 3B). In CA3, we similarly identified a significant 4-way interaction between Cluster, Diet, LPS and Sex. Saline treated female mice showed increased proportions of ameboid and decreased proportions of hypertrophic microglia in response to LPS. Both butyrate and propionate significantly blunted the LPS-induced increase in ameboid microglia, while acetate showed a similar trend but did not reach significance (Table S5). In comparison, male mice had increased ameboid microglial proportions after LPS treatment, which were was not altered by SCFA treatment.

In the PFC and DG, there were significant effects of Cluster by LPS injection and significant Sex effects; however, SCFAs did not modulate the LPS induced changes (Fig S6A-B, Table S5). Interestingly, though many genes involved in cell cycle regulation were highlighted in the transcriptomic analysis, there were no significant impacts of SCFA administration on microglial density (Fig S6C, Table S5), suggesting that microglial replication and density may not be directly correlated in a brain environment. Overall, these findings indicate that SCFAs blunt LPS-induced transitions toward an ameboid microglial morphology in a sex- and metabolite- dependent manner.

### 3.2 *In vitro* SCFA treatment alters microglial responses to LPS administration

To model the effect of SCFA administration on LPS responses *in vitro,* we treated BV2 mouse immortalized microglial cells with 250 μM of each SCFA or PBS for 1 hour, then added 10ng/mL LPS or vehicle (Fig 4A). Cells were collected after 24 hrs for downstream functional analysis.

**Figure 4.**
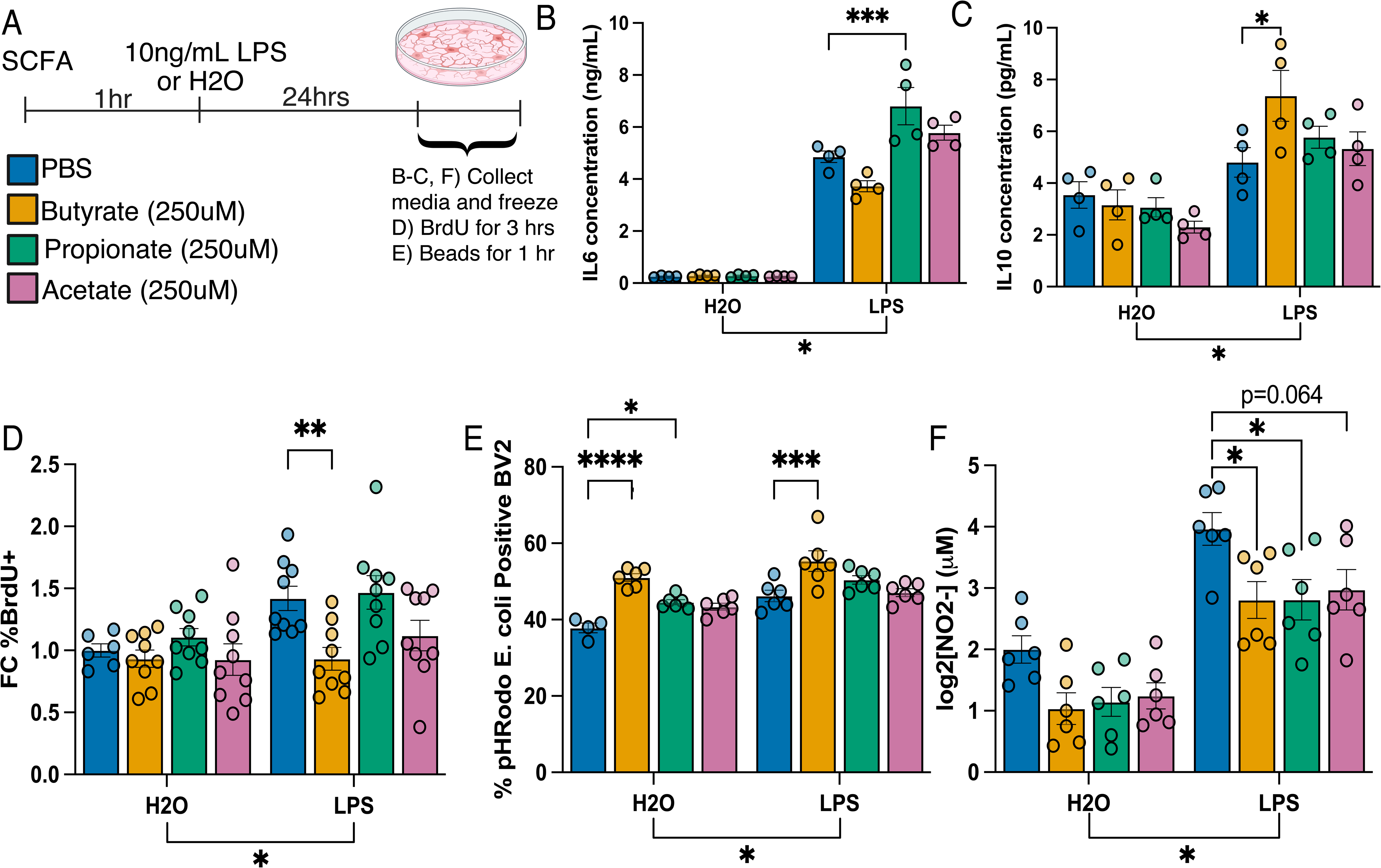
SCFAs modulate microglial inflammation response *in vitro*. (A) Outline of SCFA and LPS treatment paradigm in BV2 cells. BV2 cells were treated with PBS, Butyrate (250uM), Propionate (250uM), or Acetate (250uM) for 1 hour. LPS (10ng/mL) or H2O was added to the wells and incubated for 24 hrs prior to collection. For proliferation experiments, BrdU was added for 3 hrs after the 24 hr treatment. For phagocytosis experiments, pH-rodo *E. coli* BioParticles were added for 1 hr after the 24 hr treatment. Figure made using biorender. (B) Multiplex Protein measurements for IL6 and (C) IL10. Shown as bar graph of concentration (unit ng/mL or pg/mL) ± SEM. ANOVA main effect of LPS denoted below the graph. Tukey’s post-hoc comparisons denoted by (* p<0.03, ** p<0.002, *** p<0.0002). n=4/condition. (D) DNA replication assessment of BV2 cells treated with SCFAs by flow cytometry measured as percentage of positive cells labelled with BrdU, gated from FMO controls followed by normalization to PBS-H_2_O vehicle controls. Mean ± SEM. Tukey’s post hoc comparisons denoted by (** *p*<0.002). n=6-9/condition. (E) Phagocytosis assay of BV2 cells quantified by flow cytometry. Measured as percentage of cells positive for engulfed beads, gated from bead-negative controls. Mean ± SEM. Tukey’s post-hoc comparisons denoted by (* *p*<0.03, *** *p*<0.0002, **** *p*<0.0001). n=4-6/condition. (F) NO release into culture media, quantified with Griess Reagent Assay. Mean ± SEM Tukey’s post-hoc comparisons denoted by (* p<0.03).

#### 3.2.1 *In vitro* SCFA administration modulates inflammatory responses

To examine if the metabolite-dependent effects we observed *in vivo* were recapitulated in an *in vitro* setting, we aimed to assess the production of inflammatory cytokines by Multiplex Assay. The *in vivo* transcriptomic data suggested that SCFA supplementation would modulate LPS-induced inflammatory responses in a metabolite and sex specific manner with stronger effects observed in females. As BV2 cells are originally derived from an early postnatal female microglia culture, we hypothesized that *in vitro* SCFA supplementation would blunt production of inflammatory cytokines.

Culture media was collected for the Multiplex Assay to measure production of IL-1β, IL-6, TNF-α, and IL-10 following administration of SCFAs followed by LPS (Fig 4B-C, FigS7A-B, Table S6). For IL-6, two-way ANOVA showed a significant main effect of LPS, SCFA administration and a significant interaction (Table S6). Interestingly, propionate significantly enhanced LPS-induced production of IL-6 (p=0.0005) while butyrate blunted the effect of LPS (p=0.0537) supporting the previously observed metabolite-dependent effects (Fig 4B). For TNF-α, two-way ANOVA revealed a main effect of LPS and SCFA but no interaction. Propionate alone increased LPS-induced TNF-α production (p=0.0055) (FigS7A). For IL-10, two-way ANOVA revealed a main effect of LPS but no significant effect of SCFA administration or an interaction (Table S6). However, Tukey corrected post-hoc analysis revealed a significant enhancement of LPS induced IL-10 production by butyrate alone (p=0.0216) (Fig4C). There were no impacts of SCFA treatment on baseline cytokine levels nor impacts on modulating IL-1β at baseline or under LPS stimulation (Fig S7B) (Table S6). Together these data support the findings of metabolite-dependent modulation of LPS responses and support an anti-inflammatory role of butyrate while propionate showed a more pro-inflammatory pattern of regulation on cytokine protein levels.

#### 3.2.2 *In vitro* SCFA administration enhances phagocytosis and blunts inflammation induced proliferation and nitric oxide production

As the in vivo transcriptomics indicated a SCFA-mediated blunting of cell cycle related genes induced by LPS administration, we hypothesized that SCFAs would similarly decrease the rate of proliferation induced by LPS *in vitro.* BrdU, a thymidine analog, was administered during the last 3 hrs of LPS incubation (Fig4D) to be incorporated into newly dividing cells (Fig S8D). Two-way ANOVA revealed a significant main effect of LPS and a significant effect of SCFA administration (Table S6). Interestingly, Tukey corrected post-hocs showed that butyrate (p=0.0065) uniquely blunted LPS induced proliferation (Fig 4D). This data agrees with the *in vivo* gene expression findings, as butyrate showed the strongest blunting of cell cycle related pathways in female mice. We also examined cell viability and found no impact of SCFAs on viability (Fig S7C).

Additionally, pathways involved in phagocytosis regulation were consistently enriched in the SCFA transcriptomic data. Thus, we hypothesized that SCFA administration would impact phagocytic behavior *in vitro.* We evaluated microglial phagocytic capability by administering pH-rodo *E. coli* BioParticles for 1 hr after the 24hrs of LPS and quantifying engulfment using flow cytometry (Fig 4E). Upon engulfment and acidification of the lysosome, the pH-rodo beads fluoresce indicating phagocytic microglia (Fig S8E). Two-way ANOVA showed significant main effects of LPS and SCFA administration on the percent of bead-positive BV2s (Table S6). Tukey’s corrected post-hoc tests revealed that butyrate and propionate induced microglial phagocytosis at baseline (p<0.0001, p=0.019). Furthermore, butyrate significantly enhanced LPS induced phagocytosis (p=0.0002), while acetate and propionate did not (p>0.05) (Fig 4E). These data support that SCFAs may enhance phagocytosis at baseline, while butyrate alone enhances phagocytosis under inflammatory conditions. This aligns well with the transcriptomic data, as pathway enrichment for phagocytosis related genes in saline treated mice occurred across SCFA treatments, but were most evident after LPS administration in butyrate treated mice.

Lastly, as Arg1+ microglia gene lists were enriched in our transcriptomic data set, we hypothesized that SCFA administration *in vitro* would alter the fate of L-arginine, driving a blunting of nitric oxide (NO) production (Fig 4H). To examine this, we employed a Griess Assay to assess NO concentration in the BV2 media 24 hrs after LPS administration (Fig 4F). Two-way ANOVA revealed a significant main effect of LPS and SCFA administration (Table S6). Tukey’s corrected post-hocs revealed that all SCFAs blunted NO release, though only butyrate and propionate reached significance (p=0.0227 (Butyrate), 0.0234 (Propionate), 0.0644 (Acetate)) (Fig 4F). Together these data support a metabolite-dependent mechanism of SCFA blunting of microglial inflammatory signalling but increased phagocytic capacity.

### 3.3 SCFAs enact modulation of inflammatory responses through regulation of histone acetylation and inhibition of HDAC activity

We next further probed the mechanism through which SCFAs modulate microglial inflammatory responses. Previous research has supported a model in which butyrate acts on gene expression by inhibiting HDAC activity in microglia (Caetano-Silva et al., 2023). Interestingly, our previous BV2 study showed that inhibition of HDAC3 promoted enhancements of *Arg1* expression resulting in functional outcomes that mirror our findings on SCFA inhibition(Meleady et al., 2023). Therefore, we hypothesized that SCFAs could act as inhibitors of HDAC activity, enhancing histone acetylation.

To examine this *in vivo*, we employed a flow cytometry-based method to assess histone acetylation on a global level *in vivo* (Towriss et al., 2023a). Twenty-four hours after LPS or VEH administration, microglia were isolated and analyzed by flow cytometry for global levels of H3K9ac and H3K27ac (Fig 5A-C, Table S7). ANOVA analysis with batch correction revealed a significant main effect of LPS for both H3K9Ac and H3K27Ac, but no significant main effect of SCFA administration nor interaction. However, Sidak post-hoc analysis revealed that SCFA supplementation enhanced H3K9Ac above the level of the saline control in LPS administered female mice (p=0.002 (butyrate), 0.003 (propionate), 0.030 (acetate)) (Fig 5B). Butyrate alone also enhanced H3K9ac levels in male vehicle administered mice (p=0.013). Additionally, H3K27ac was enhanced by SCFA supplementation in LPS administered females (p=0.0002 (butyrate), 0.004 (propionate), 0.011 (acetate) (Fig 5C). Butyrate alone again enhanced H3K27ac levels at baseline in male mice (p=0.013).

**Figure 5.**
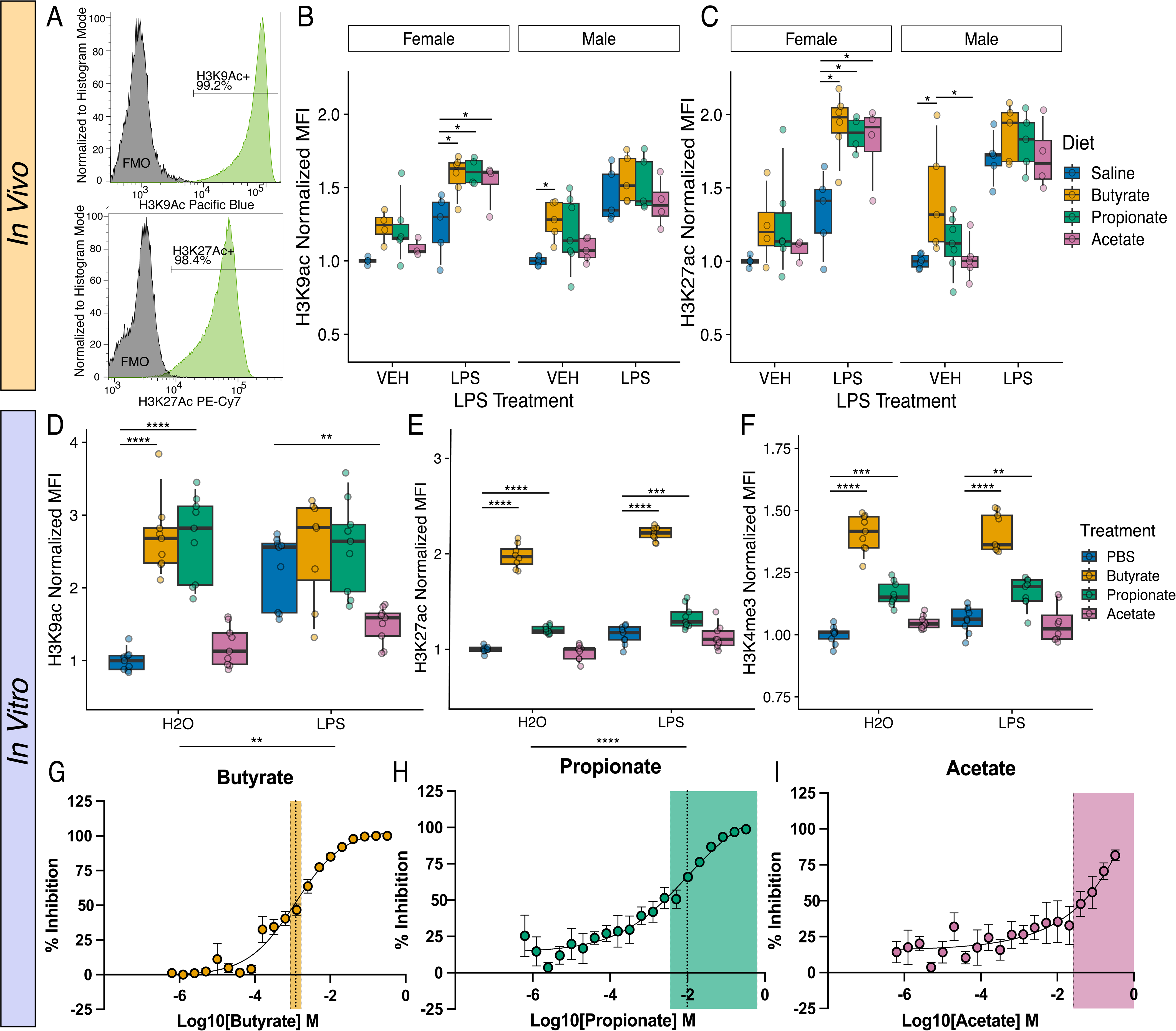
Butyrate acts as an HDAC inhibitor resulting in enhanced global histone acetylation at baseline and in response to LPS. (A) Isolated microglia cells (CD11b+CD45+) were assessed for signal on either H3K9ac-PacBlue or H3K27ac-PECy7. (B) The median fluorescence intensity (MFI) for total levels of H3K9ac and (C) H3K27ac was normalized to MFI of vehicle controls within each sex (* *p*<0.05). n=4-6/condition. (D) In BV2 cells, H3K9ac, (E) H3K27ac, and (F) H3K4me3 MFI was assessed following SCFA and LPS treatment. MFI for total levels of histone modifications in all BV2 samples were normalized to MFI of PBS-H_2_O controls. Tukey’s post-hoc comparisons denoted by (** *p*<0.002, *** *p*<0.0002, **** *p*<0.0001). (G) Percent inhibition of HDAC Activity [(RLU(Sample)-RLU(Background))/(RLU(PBS)*100] was evaluated for butyrate, (H) propionate and (I) acetate in BV2 cells. Data were fitted to a non-linear, sigmoidal, four-parameter logistic regression. Data exceeding 100 or 0 was removed to the respective value for regression stability. IC50 ± 95%CI is depicted for each metabolite as shaded bars.

To examine if this pattern of changes replicated *in vitro*, we employed the same SCFA+LPS BV2 model to assess the global levels of H3K9ac, H3K27ac, and H3K4me3, which are all markers of open chromatin associated with histone hyperacetylation (Fig 5D-F). Two-way ANOVA revealed LPS and SCFAs had a significant effect on H3K9ac, H3K27ac and H3K4me3 (Table S7). LPS enhanced all three histone marks across SCFA conditions. Sidak post-hoc comparisons demonstrated significant enhancement of total H3K27ac for butyrate, at baseline (p<0.0001) and in response to LPS (p<0.0001) and propionate, at baseline (p=0.0001) and in response to LPS (p=0.0004). Similarly, H3K4me3 was significantly enhanced by butyrate, both at baseline (p<0.0001) and in response to LPS (p= p<0.0001), and propionate, both at baseline (p<0.0001) and in response to LPS (p=0.001). Acetate had no effect on either H3K27ac or H3K4me3 (Fig 5E-F). H3K9Ac was significantly increased at baseline both by butyrate (p<0.0001) and propionate (p<0.0001); however, not under LPS stimulation, while acetate suppressed LPS induced H3K9ac levels (p=0.009)(Fig 5D). Together, these *in vitro* data support the *in vivo* findings that butyrate has the strongest impact on microglial histone acetylation, which is consistent with the strongest impact on microglia transcription and downstream functions.

Finally, as we found global shifts in histone acetylation and the strongest role of butyrate in mediating SCFA impacts on microglia, we hypothesized that SCFAs may impact microglia by inhibiting HDAC activity. To examine this inhibition directly, we performed a dilution curve series using a luciferase-based assay to measure HDAC inhibition by SCFA (Fig 5G-I, Table S7). In this assay, the HDAC-Glo^TM^ I/II substrate is deacetylated by HDACs and, upon addition of the developer reagent, creates the luminescent compound aminoluciferin. Resultingly, *in vitro* inhibition of HDAC activity by SCFAs would reduce the ability of HDAC to deacetylate the substrate resulting in lower fluorescence. BV2 cells were a dose curve of with SCFAs ranging from 0.3125 μM to 327.68 mM with Trichostatin A as a positive control (Fig S7D). The concentration at which 50% of HDAC present was inhibited (IC50) for butyrate was determined to be 1.2 mM (0.869mM-1.690mM, R^2^ = 0.9499) (Fig 5G). While the IC50 for propionate was determined to be 9.8mM (3.5mM-638.1mM, R^2^ = 0.8346) and acetate was determined to be 32148M (260mM-DNE, R^2^ = 0.4899) (Fig 5H-I). Notably, the upper limit of propionate and acetate IC50 were outside of the range of testing as the upper plateau was not confidently determinable. This suggests that while butyrate concretely has significant HDAC inhibition activity, we could not confidently determine that propionate and acetate possessed HDAC inhibition capabilities. This is consistent with the strongest enhancement of histone acetylation by BV2 cells *in vitro*. Together these data suggest that SCFA modulation of microglia is sex- and metabolite- specific and functions through remodelling of the epigenomic landscape by inhibition of HDAC activity, specifically by butyrate and to a lesser extent propionate and acetate.

## 4. Discussion

Emerging evidence suggests that metabolites derived from gut microbiota may be critical regulators of microglial function and may influence the process of inflammation in the CNS (Erny et al., 2015; Spielman et al., 2018; Thion et al., 2018). While disruption of SCFA signalling has been shown to cause microglial dysfunction, the mechanism by which SCFAs regulate microglial behavior remains unclear. In the present study, we demonstrated that SCFA supplementation through drinking water modulates microglial inflammatory responses in a sex- and metabolite-specific manner. Specifically, SCFA treatment reduced LPS-driven sickness behavior, blunted inflammatory gene expression and morphology changes and enhanced microglial histone acetylation levels, potentially leading to shifts towards more homeostatic programming during inflammatory challenge. However, further studies are needed to clarify how SCFA treatment influences chromatin organization and histone acetylation at specific genomic loci. Interestingly, of the three SCFAs that we evaluated, butyrate had the most significant impact on the inflammatory response in LPS-administered female mice. When we moved into BV2 microglia-like cells which are an immortalized female microglial line, we observed a similar impact of butyrate on inflammatory programming. Here, we demonstrated that *in vitro* butyrate acts as a potent HDAC inhibitor, increasing global histone acetylation levels and altering the production of inflammatory cytokines. We further showed that butyrate promoted a more homeostatic phenotype in response to inflammation by blunting proliferation and NO production while enhancing phagocytosis. These results correlated well with the *in vivo* transcriptomic findings, which suggested enhancements in phagocytic and Arginase 1 positive microglia associated gene programming. Altogether, our data support a model in which SCFAs produce immunomodulatory effects by promoting a more homeostatic and neuroprotective microglial state.

Previous studies have shown that SCFAs can induce immunomodulatory effects by inhibiting HDAC activity and stimulating histone acetyltransferases resulting in enhanced histone acetylation (Dalile et al., 2019; Mann et al., 2024; Thomas and Denu, 2021). Specifically, several studies have previously shown that SCFAs can inhibit HDAC activity *in vitro* (Caetano-Silva et al., 2023; Jansen et al., 2004; Waldecker et al., 2008). Here, we similarly observed global increases in histone acetylation upon SCFA treatment both *in vitro* and *in vivo,* with butyrate causing the most significant increase. Butyrate has previously been shown to inhibit Class I and II HDACs by interfering with zinc-dependent catalytic activity (Davie, 2003; Reddy et al., 2018; Steliou et al., 2012). In BV2 cells, butyrate may act more specifically through inhibition of HDAC3, as we observed that the impact of butyrate on the LPS response was remarkably similar to that which we observed in our previous study utilizing RGFP966, an HDAC-3 specific inhibitor (Meleady et al., 2023). One example of this similarity is the blunting of LPS-induced NO production which is affiliated with enhanced Arg1+ microglial gene regulation and Arg1 gene expression (Meleady et al., 2023; Stratoulias et al., 2023). Arginase competes with iNOS for a shared substrate (L-arginine), as such Arg1+ microglia may be less prone to NO production and subsequent inflammation. Both our *in vivo* transcriptomic data and *in vitro* functional data support that butyrate administration blunts LPS-induced NO production by increasing genes involved in the arginase pathway. Excessive NO production can impair homeostatic microglial metabolism by inhibiting mitochondrial respiration, limiting oxidative metabolism and contributing to enhanced glycolysis. Metabolic reprogramming to favour glycolysis is an essential part of the microglial inflammatory response (Bernier et al., 2020; Holland et al., 2018; Orihuela et al., 2016; Towriss et al., 2023b). As such, a reduction in the NO production observed *in vitro* may reflect broader alterations in inflammatory metabolic reprogramming supporting the overall blunting of inflammatory responses that we observed *in vivo*. Additionally, previous work by Cherry *et al*. and Stratoulias *et al*. support that Arg1+ expression in microglia promotes expression of genes linked to a unique subtype of microglia that may be highly phagocytic and are associated with a reduction in plaque deposition in an AD model (Cherry et al., 2015; Stratoulias et al., 2023). Similarly, our data also supported that SCFA administration *in vitro* was associated with enhanced phagocytosis, supporting the model that metabolite administration through HDAC inhibition may promote Arg1+ microglial traits, including enhancing phagocytosis. This interpretation aligns with previous studies demonstrating that HDAC3 contributes to inflammatory reprogramming in myeloid cells (Chen et al., 2012; Mullican et al., 2011; Rosete and Ciernia, 2024). Together these findings support a hypothesis that butyrate may operate through HDAC3 specific inhibition, which would be an interesting avenue for future research.

In addition to neuroimmune modulation, SCFAs have been implicated in behavioral regulation (Tang et al., 2022; Van De Wouw et al., 2018). We found that SCFA administration alone did not alter responses to behavior tests; however, SCFAs did blunt LPS-driven sickness behaviors. Although similar SCFA modulation of sickness has not been reported previously, butyrate has been shown to reverse LPS-induced depressive-like and stress-induced anxiety-like behaviors by acting in an anti-inflammatory role (Van De Wouw et al., 2018; Yamawaki et al., 2018). Likewise, outside of the CNS, butyrate has been suggested to be pro-homeostatic in other tissue resident macrophages, resulting in amelioration of inflammation in alcoholic liver disease, obesity, and septic shock (Rafiei et al., 2024; Ren et al., 2022; Wang et al., 2017). Interestingly, although LPS-induced microglial inflammation can contribute to sickness behavior, microglia are not essential, as their depletion does not abolish LPS-driven sickness (Vichaya et al., 2020). Thus, sickness repression driven by butyrate is likely to involve additional peripheral immune regulation.

One major limitation of our study is that our *in vitro* system was unable to examine the impact of sex on metabolite responses as BV2 cells are derived from a female post-natal mouse microglial culture. Huuskonen *et al*. found contrasting effects to those we observed *in vitro* while utilizing the N9 cell line, which is derived from a male post-natal mouse microglial culture (Huuskonen et al., 2004). Ultimately supporting that sex-specific effects may persist *in vitro* agreeing with the sex-specific effects our in vivo findings. Additionally, previous *ex vivo* studies with primary microglial cultures have observed contrasting effects which may arise from failure to include sex as a variable in their studies (Caetano-Silva et al., 2023; Churchward et al., 2023; Colombo et al., 2021). Interestingly, de Witte *et al*. examined sex-specific effects of butyrate administration in microglial cultures and similarly found that female cultures exhibited a stronger response to butyrate when compared to male cultures (Vichaya et al., 2020). As such our findings are in agreement with a large body of literature documenting significant sex-effects of microglial responses that exist both *in vitro* and *in vivo* (Sullivan and Ciernia, 2022; Thion et al., 2018). Given that there are significant documented sex-effects linked to disease-associated pathways and disease-risk, these data add to the literature supporting the importance of thoroughly assessing sex differences in the development and assessment of therapeutics (Eid et al., 2019; McCarthy and Wright, 2017; Sullivan and Ciernia, 2022).

Similarly, a limitation of our study is that SCFA supplementation was initiated in adulthood (>8 weeks), potentially overlooking developmental periods during which microbial metabolites are known to shape microglial maturation (Erny et al., 2021; Silva et al., 2020; Thion et al., 2018). Early life microbial colonization and SCFA availability have been shown to be critical in establishing normal microglial identity and immune function, as such supplementation in development may have more profound impacts. Consequently, our findings can only be interpreted as reflecting the effects of SCFA supplementation on mature microglia and, therefore, are more relevant to evaluating the potential of SCFA supplementation as a therapeutic strategy in adulthood rather than the role in developmental programming. Finally, another limitation of the study is that the concentration of metabolites examined *in vitro* may not reflect physiologically relevant concentrations in the brain microenvironment. While the concentrations used in this study were selected based on previous literature and aimed to reflect circulating concentrations in murine serum, the concentrations of SCFA in the brain are likely considerably lower what was examined in this study (Cummings et al., 1987; Huuskonen et al., 2004; Silva et al., 2020; Sun et al., 2016; Zeng et al., 2022). Consequently, the direct effects observed *in vitro* may not recapitulate *in vivo* exposure where microglia are subjected to lower metabolite concentrations and more diverse environmental signals. Future studies should more directly quantify physiological and pathological concentrations of SCFAs in the brain microenvironment both in homeostasis and in response to oral SCFA supplementation to establish a dose-response relationship and more closely bridge *in vitro* and *in vivo* models of SCFA impacts on microglia.

Our data supports that SCFAs have sex- and metabolite-specific immunomodulatory effects and that butyrate may promote homeostatic programming ultimately resulting in a neuroprotective phenotype. These findings are important as they highlight the necessity of investigating SCFAs as individual metabolites that may act through unique pathways to enact their functions. Our study suggested that butyrate was the most significant regulator of HDAC function and subsequent neuroinflammation. Importantly, as we administered SCFAs to mice with intact microbiomes, endogenous SCFA production may have limited the magnitude of the observed *in vivo* impacts through potential ceiling effects. This interpretation is consistent with previous literature demonstrating that probiotic administration often produces modest phenotypic changes in healthy individuals (Éliás et al., 2026; Kristensen et al., 2016; McFarland et al., 2018). However, as disruption of SCFA-producing bacterial populations has been associated with microglial dysfunction, we hypothesize that supplementation of SCFAs in these models may have a more prominent effect in ameliorating disease phenotypes (Spielman et al., 2018; Sullivan et al., 2025). Similarly, as the animals had commensal microbiomes, there could be microbiota-mediated effects *in vivo* that would not be recapitulated *in vitro.* For example, oral supplementation of SCFAs may change the gut microbiota composition causing changes in metabolite presence (Huang et al., 2023) or change animal appetite resulting in changes that would not be directly related to metabolite changes (Li et al., 2018). While our behavior studies and animal monitoring suggest no change in animal weight or water consumption because of metabolite administration, we did not directly examine any non-specific effects of SCFA supplementation.

Our work highlights the potential of SCFA supplementation as postbiotic candidates for neurological disease treatment in the context of microglial dysfunction. Our findings show that oral SCFAs are well tolerated in vivo and blunt sickness behavior without altering normal locomotion or anxiety-like behavior, suggesting they may serve as a powerful postbiotic for disease treatment. Gut dysbiosis linked to deficiency of SCFA-producing microbes has been previously linked to neurodevelopmental (Kang et al., 2013; Parracho et al., 2005; Vuong and Hsiao, 2017), neuropsychiatric (Jiang et al., 2015; Painold et al., 2019; Xu et al., 2012), neurodegenerative diseases including AD (Cattaneo et al., 2017; Hill et al., 2014; Vogt et al., 2017) and PD (Hill□Burns et al., 2017; Scheperjans et al., 2015). Interestingly, previous research has shown that fecal microbiota transplant (FMT) can cause significant improvements for neurological disorders (Chen et al., 2025; Churchward et al., 2023; Eslami et al., 2025; Park et al., 2022; Zhang et al., 2023). Strikingly, previous research has shown improvement in these conditions with SCFA supplementation alone (Erny et al., 2021; Mezö et al., 2020). As such, future directions for this work could look more directly at how specific clinically-relevant dysbiosis contributes to microglial dysfunction to highlight the potential of SCFA supplementation and HDAC-activity manipulation in improving neurological symptoms.

Collectively, our findings support the therapeutic potential of gut-derived metabolites, particularly butyrate, in the context of neurological disorders associated with dysbiosis. Butyrate appears to function as an HDAC inhibitor, promoting epigenomic reprogramming that favors protective transcriptional responses during inflammation. Notably, our results suggest that butyrate-mediated HDAC inhibition reduces inflammatory signalling while enhancing microglial phagocytosis, thereby facilitating pathogen clearance and debris removal, increasing the resolution of inflammation. Overall, this work establishes a foundation for future investigations into brain disorders linked to gut dysbiosis, highlighting how distinct SCFAs, including butyrate, propionate, and acetate, modulate microglial function.

## Declarations

## Acknowledgements

We are grateful for the feedback and support from every member of the Ciernia Lab throughout the project. We want to especially thank Jia Gandhi and Kaitlyn Tonary for their help with changing water treatments and weighing mice for the *in vivo* experiments. We want to sincerely thank the animal care staff, Rena Chen, Kathen Li, Emily Yeh and many others, for all their work in caring for the animals in this study and consistently monitoring the water levels in these cages. We also thank Eve Technologies for their guidance and prompt responses regarding the Multiplex Protein Assay. We are grateful for the resources provided by the Neuroimaging & Neurocomputation Centre and the Dynamic Brain Circuits in Health & Disease Research Cluster at the UBC Djavad Mowafaghian Centre for Brain Health for computational resources (RRID SCR_019086).

## Ethics approval

All animal experiments were approved by the Canadian Council on Animal Care, and all protocols were approved by the Animal Care Committee of the University of British Columbia (A23-0086).

## Availability of data and materials

RNA-seq data is available at NCBI GEO at GSE306136. All analysis code is available at https://github.com/ciernialab/Towriss-et-al.-2026

## CRediT Author Contributions

**Conceptualization**: A.V.C., V.D., M.T., and C.W. conceptualized the project. **Methodology/Investigation**: M.T. performed most in vivo experiments including animal maintenance (colony preparation, water changes, monitoring, injections), microglia isolations, FACS sorting and IHC and assisted with bioinformatic analysis. V.D. performed most of the in vitro experiments, microglia isolations, qFLOW experiments. J.G assisted with in vivo experiments and performed the in vitro HDAC inhibitor assay experiments. J.C performed behavior experiments and assisted with microglia isolations. C.A. and K.M assisted with IHC collection and analysis. J.H, A.U and J.M.S contributed to aiding with colony water maintenance and microglia isolations. **Analysis**: M.T., V.D. and J.G. performed experiment analyses. A.V.C performed bioinformatic analysis of RNA sequencing data. **Writing original draft**: : Manuscript was written by M.T., V.D., and A.V.C. **Writing – review & editing**: All authors read, edited and approved the manuscript. **Funding acquisition & project administration:** A.V.C and C.W.

---

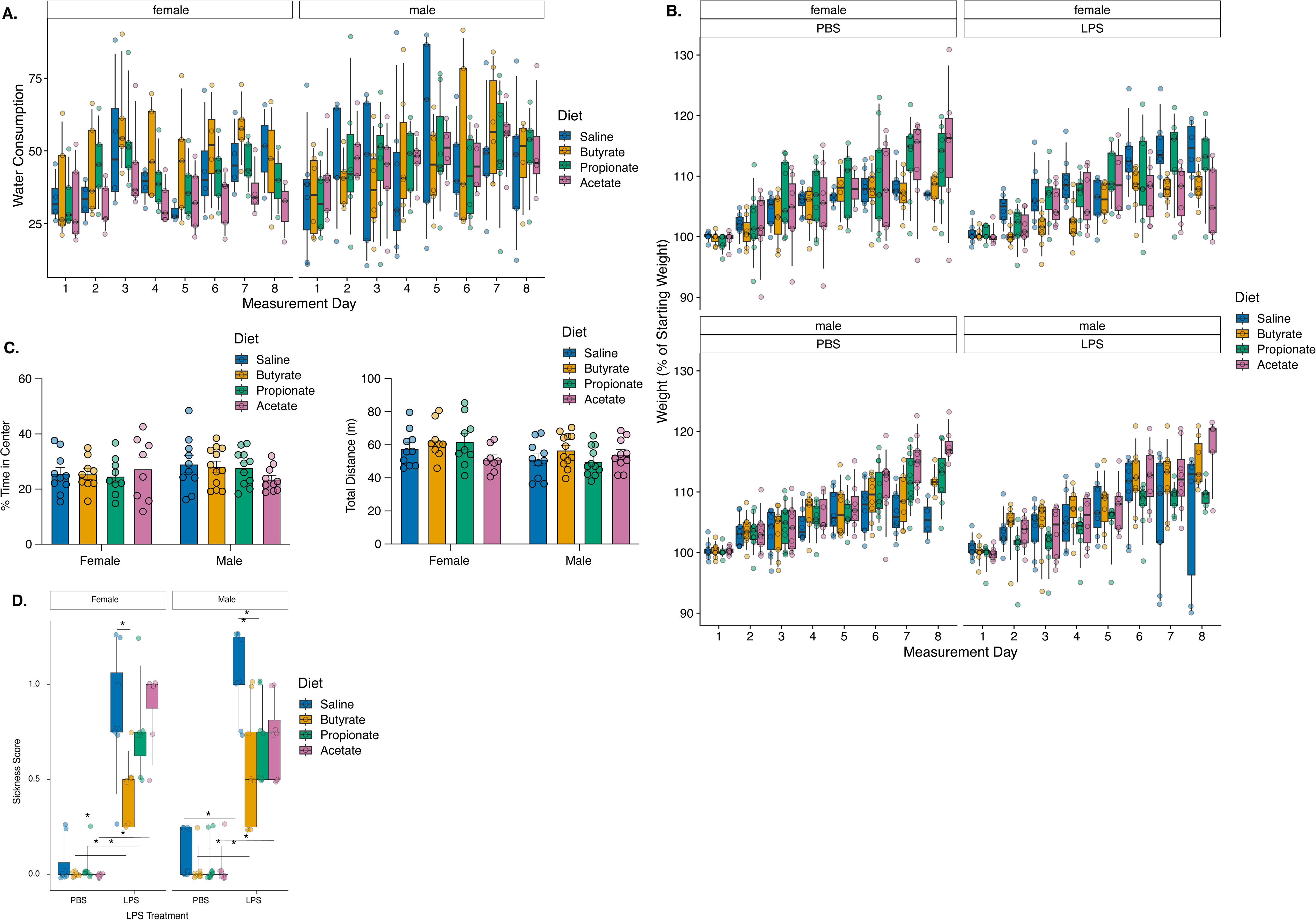

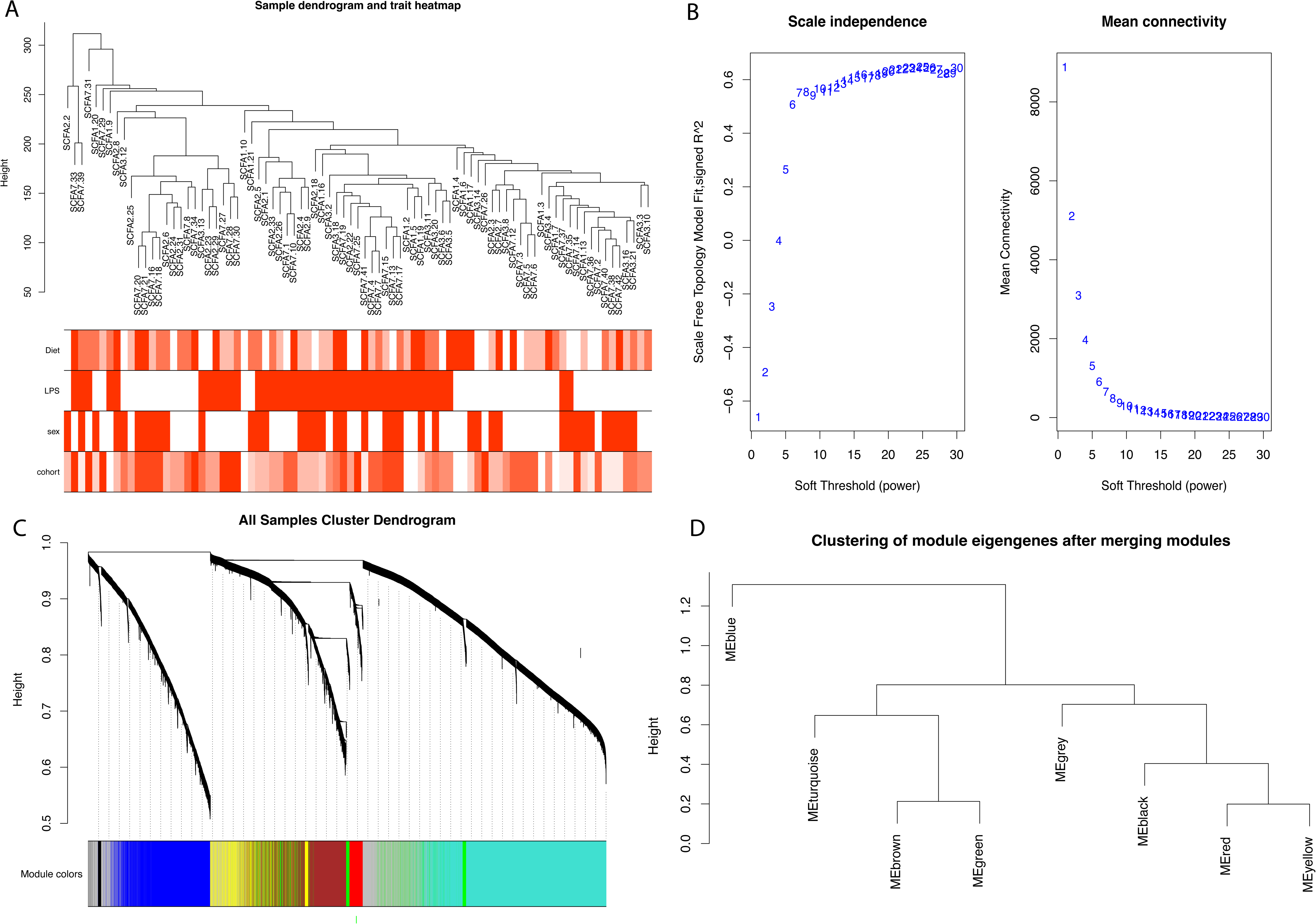

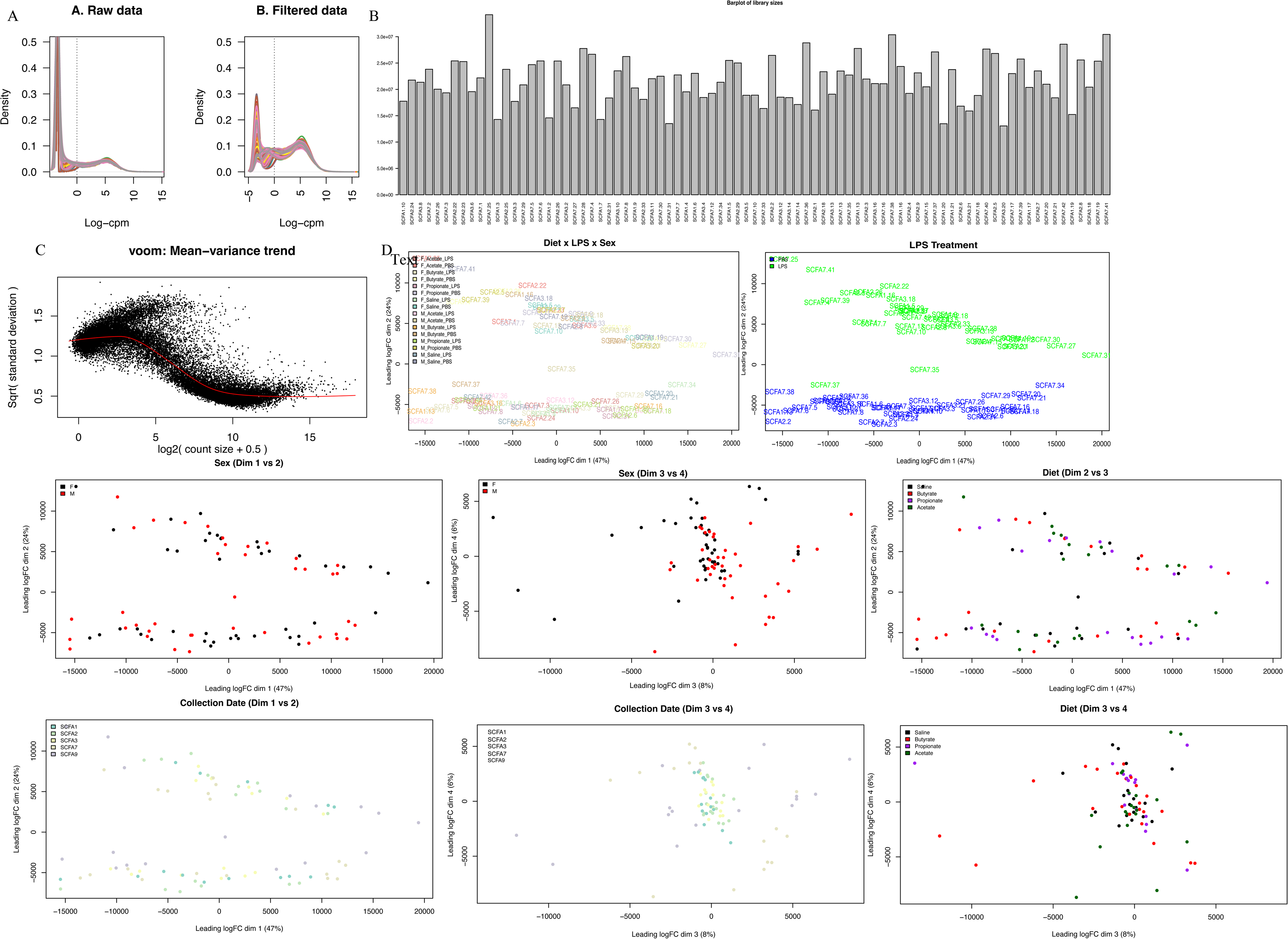

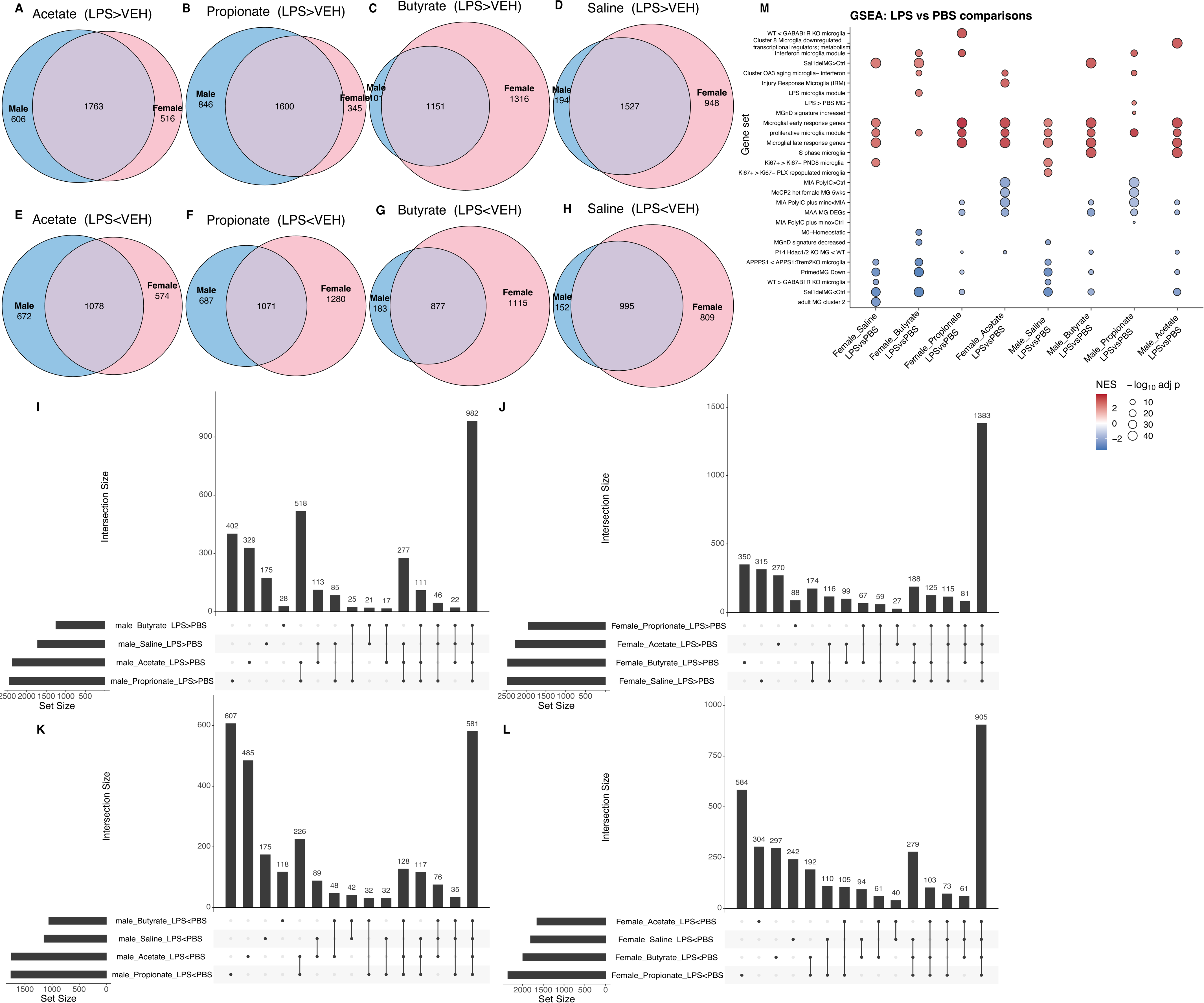

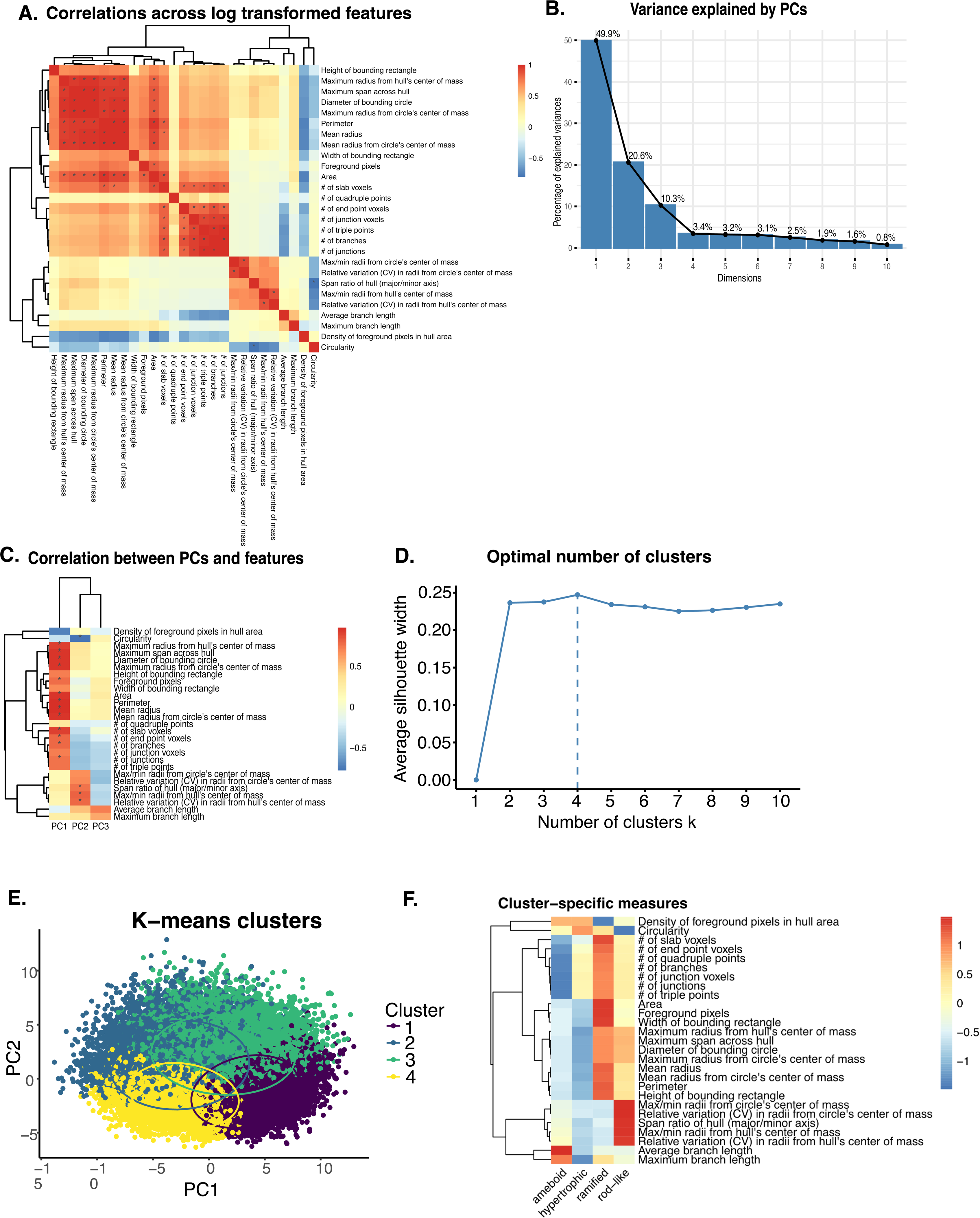

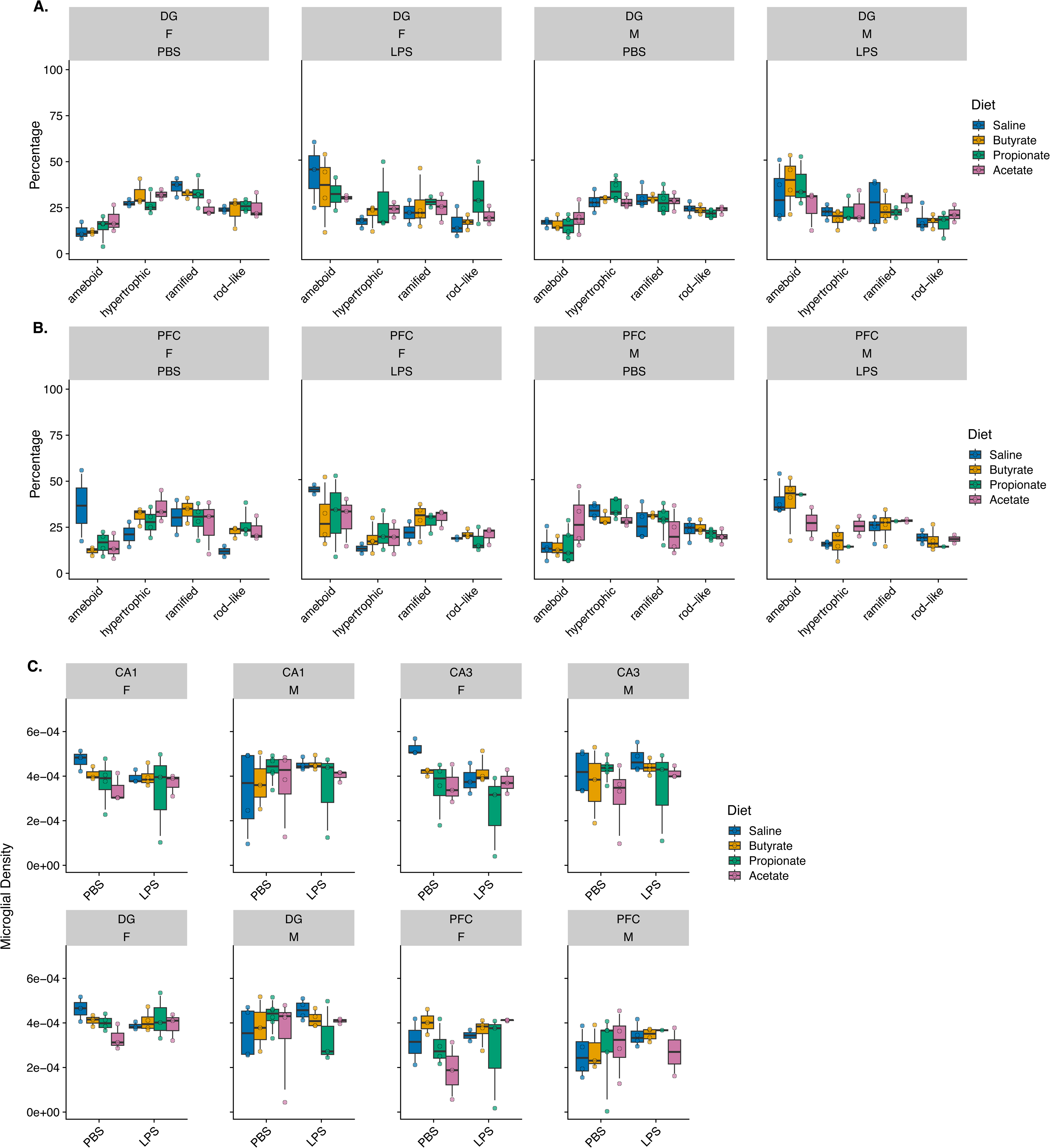

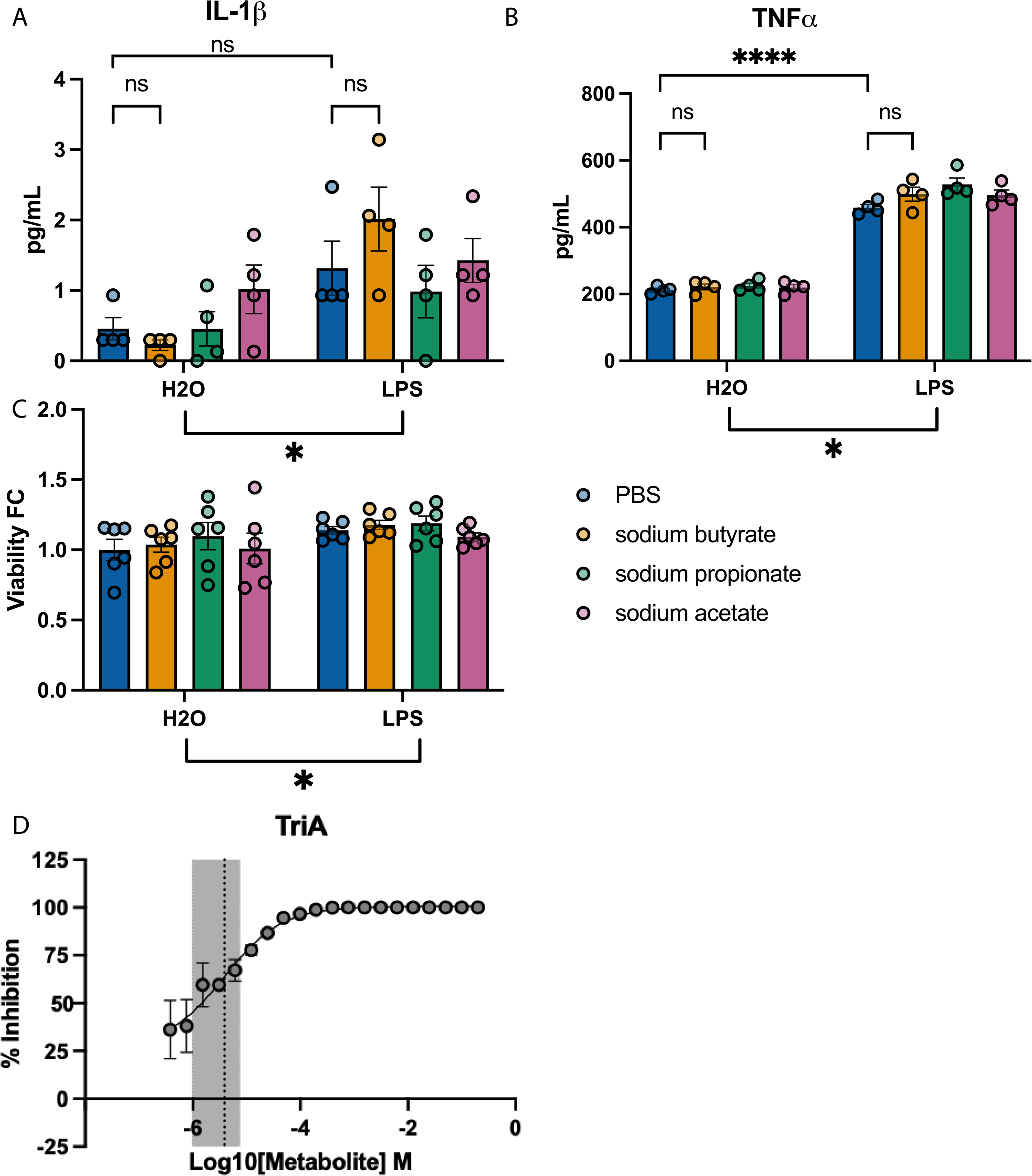

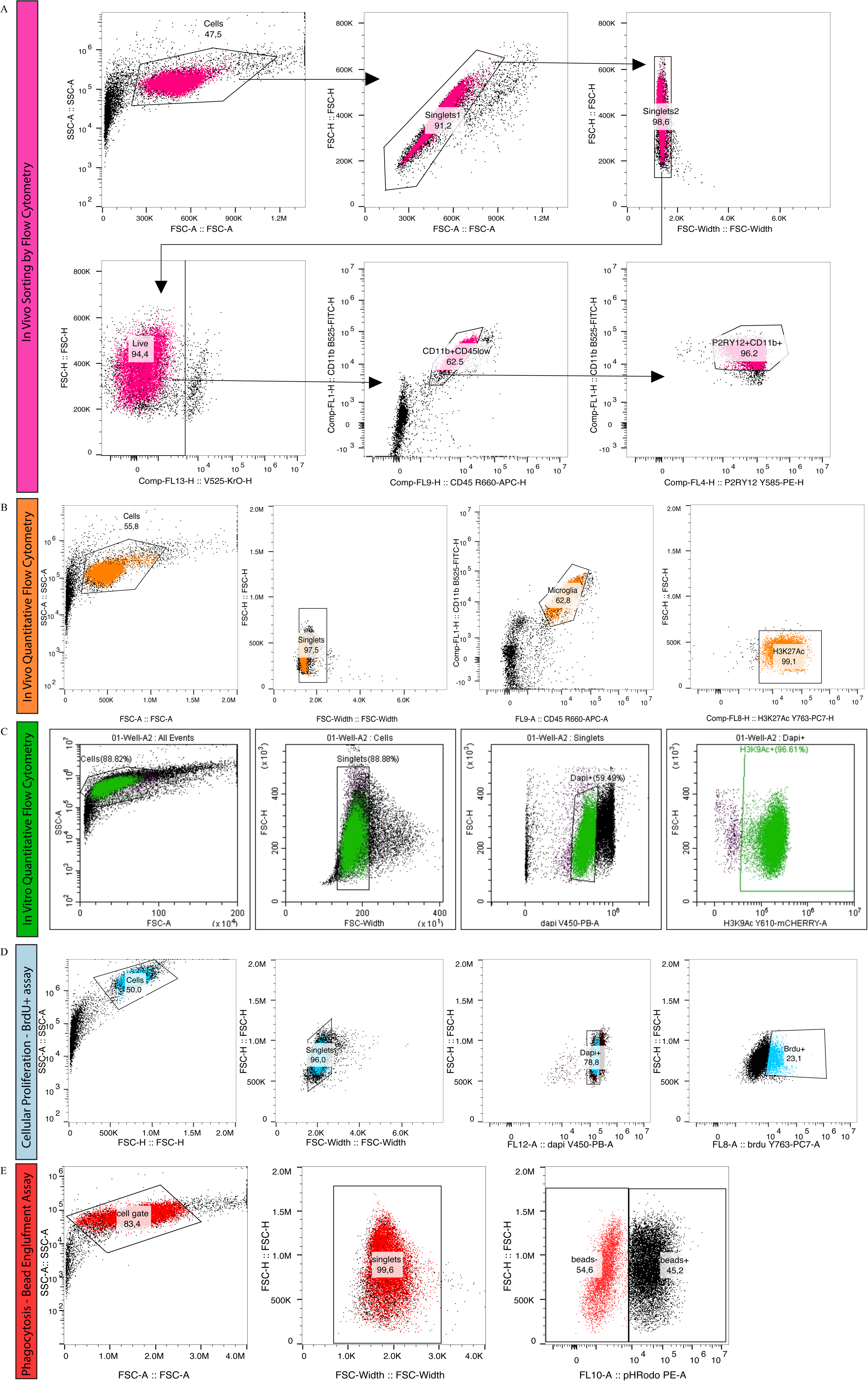

