## Supplement Captions for "Microbiome-derived Short Chain Fatty Acids modulate microglial inflammatory responses in a sex- and metabolite-specific manner"

**Supplemental Legends**

**Supplemental Figures**

Figure S1. **Physiological and behavioral effects of SCFA treatment.**
A. Water consumption measured across repeated time points and plotted by diet condition (Saline, Butyrate, Propionate, Acetate), shown separately for females and males. Measurement days correspond to sequential collection intervals. Boxplots depict the median and interquartile range, with whiskers indicating the 5th–95th percentiles. B. Body weight expressed as a percentage of starting weight over time, stratified by sex and injection condition (PBS or LPS). Data are plotted across measurement days and colored by diet. Boxplots depict the median and interquartile range, with whiskers indicating the 5th–95th percentiles. C. Open‑field behavioral outcomes, including percent time spent in the center and total distance traveled, shown for females and males across dietary conditions. Mean +/- SEM. Individual animals shown as data points. D. Sickness scoring (0-3) 3 hours after LPS or PBS injection. LPS significantly increased sickness in all Diet conditions for both sexes. Butyrate significantly blunted sickness in females and both Butyrate and Propionate blunted sickness in males. Boxplots depict the median and interquartile range, with whiskers indicating the 5th–95th percentiles. * p<0.05 Holm corrected posthocs.

Figure S2. WGCNA. A. hierarchical clustering of samples. B. Scale‑free topology analysis was performed to determine an appropriate soft‑thresholding power (β) for weighted gene co‑expression network construction. **Left:** Scale‑free topology model fit (signed R²) as a function of soft‑threshold power. **Right:** Mean connectivity plotted against soft‑threshold power, demonstrating progressive loss of network connectivity at higher powers. C. Hierarchical clustering of genes based on topological overlap is shown for all samples following weighted gene co‑expression network construction. The dendrogram was generated using average linkage clustering, and modules are indicated by color bars beneath the dendrogram. D. Hierarchical clustering of module eigengenes (MEs). Dissimilarity was calculated as 1 − Pearson correlation between module eigengenes and clustered using average linkage.

Figure S3. RNAseq Differential Analysis. A. **Expression filtering using log‑CPM distributions.** Density plots of log2 counts per million (CPM) across all samples shown before (left) and after (right) expression filtering. Vertical dashed lines indicate log‑CPM thresholds corresponding to minimal expression cutoffs. B. **Library size distribution across samples.** Bar plot showing total read counts (library sizes) for each RNA‑seq sample. **C. Mean–variance relationship modeled by voom.** Voom mean–variance plot illustrating the relationship between log2 expression level and the square root of the estimated variance following normalization. The fitted trend captures the decreasing variance with increasing expression and was used to generate observation‑level precision weights for linear modeling and differential expression analysis. D. Multidimensional scaling (MDS) plots were generated using normalized log‑CPM values for the top 500 most variably expressed genes following batch correction for experimental cohort. Percent variance explained by each dimension is indicated on the corresponding axes.

Figure S4. **Sex‑specific + Diet‑dependent overlap of LPS‑responsive genes.**
(A-H) Euler diagrams illustrate the overlap of differentially expressed genes (DEGs) between male and female microglia following acute LPS challenge (LPS vs VEH) under each dietary condition. Diagrams depict sex‑shared and sex‑specific gene sets, with numbers indicating the total DEGs unique to each sex or shared between sexes. **(A–D) LPS‑increased genes (LPS > VEH).** Euler plots showing overlap of genes significantly increased by LPS in male and female microglia under **A.** Acetate, **B.** Propionate, **C.** Butyrate, and **D.** Saline diets. **(E–H) LPS‑decreased genes (LPS < VEH).** Euler plots showing overlap of genes significantly decreased by LPS in male and female microglia under **E.** Acetate, **F.** Propionate, **G.** Butyrate, and **H.** Saline diets. (I-L) UpSet plots summarize the overlap of DEGs following acute LPS challenge (LPS vs PBS) across dietary conditions. Each set corresponds to a specific diet, and intersection bars indicate the number of genes shared among one or more diets. Horizontal bars represent the total number of DEGs identified within each dietary condition. **I. LPS‑increased genes (LPS > PBS) in males.** **J. LPS‑increased genes (LPS > PBS) in females. K. LPS‑decreased genes (LPS < PBS) in males.** **L. LPS‑decreased genes (LPS < PBS) in females.** (M) Dot plot showing normalized enrichment scores (NES) from GSEA comparing **LPS versus PBS** conditions across sex (female [F], male [M]) and treatment groups (saline, butyrate, propionate, acetate). Each row represents a curated microglial gene set from MGEnrichment, and each column corresponds to a specified comparison. Dot color indicates the direction and magnitude of enrichment (NES), with positive values reflecting enrichment in LPS relative to PBS and negative values indicating enrichment in PBS compared to LPS. Dot size represents statistical significance (−log10 adjusted p value).

**Figure S5. Analysis of microglial morphological features.**
A. Heatmap showing pairwise correlations across morphological features extracted from microglial reconstructions, including size, shape, and branching metrics. B. Percentage of variance explained by each principal component (PC), with the first three PCs accounting for the largest proportion of total variance. C. Correlation matrix illustrating the contribution of individual morphological features to PC1–PC3, highlighting features associated with cell size, ramification, and structural complexity. D. Average silhouette width across k‑means solutions, identifying the 4 optimal number of clusters used for downstream analysis. E. Projection of individual cells in PC space (PC1 vs PC2), colored by 4 k‑means cluster assignment, revealing distinct morphological states. F. Correlation between each k-means cluster and individual morphology metrics used to ascribe morphological names to each cluster.

Figure S6. Morphology and Density Metrics. A. Morphology results for dentate gyrus (DG) of the hippocampus across sexes, Diets and LPS injection conditions. Significant main effect of Cluster and Cluster x LPS interaction. No main effects of Sex or Diet and no interactions. B. Morphology results for the prefrontal cortex (PFC) across sexes, Diets and LPS injection conditions. There was a significant main effect of Cluster, a significant Cluster x LPS interaction and a significant Cluster x Diet x Sex interaction. D. Microglial density for subregions of the hippocampus (CA1, CA3, and DG) and prefrontal cortex (PFC). There were no significant ANOVA effects beyond a main effect of Diet in CA3.

Figure S7. *In Vitro* Supplement in BV2 cells. A-B. Multiplex protein assay recordings for IL-1B and TNFa. ANOVA main effect of LPS denoted below the graph. Mean ± SEM. Tukey’s post-hoc comparisons denoted by (*** p<0.0002). n=4/condition. C. XTT assay performed on 10k BV2 cells after 24 hrs treatment. Shown as a fold change to PBS H2O. Mean ± SEM. n=4/condition. D. Trichostatin A positive control for HDAC3 Glo Assay. Percent inhibition of HDAC Activity [(RLU(Sample)-RLU(Background))/(RLU(PBS)*100]. Data were fitted to a non-linear, sigmoidal, four-parameter logistic regression. IC50 ± 95%CI is depicted as shaded bars with dotted line on the IC50.

Figure S8. Gating strategies. A. Flow Sorting Gating strategy. Microglia were identified as live singlet cells that were CD11b+ CD45low and P2RY12+. B. Intracellular flow staining *in vivo.* CD11b+CD45low singlet cells (microglia) were analyzed for H3K27Ac or H3K9Ac signal on their respective channels. C. Intracellular flow staining *in vitro.* Singlet, non-dividing BV2 cells were identified by DAPI positive and H3K27Ac, H3K9Ac or H3K4me1 were evaluated on their respective channels. D. Newly divided BV2 cells were identified as DAPI positive single cells above the isotype control for BrdU-PECy7. E. Cells positive for engulfed beads were identified as singlet cells positive for pHRodo signal on PE.

**Supplemental Tables**

S1. **Physiological and behavioral effects of SCFA treatment.**

**Water Consumption: ANOVA results for liner modeling of Diet, Sex and Day of water consumption (4 day average by cage) with Cohort as a covariate and Cage as a repeated measure. Tukey’s corrected posthocs comparing the slope of change over time between Diets for within each Sex.**

**Body Weight: ANOVA results for liner modeling of Diet, Sex, LPS injection and Day with Cohort as a covariate and Animal as a repeated measure. Tukey’s corrected posthocs comparing the slope of change over time between Diets for within each Sex and LPS Treatment. Final weight collected before LPS injection.**

**Open Field: ANOVA results for % of time in the center and total distanced travelled in the open field for Sex, Diet and the Interaction.**

**Sickness Behaviour: ART analysis of sickness rating (0-3) and Holm corrected posthocs.**

S2. WGCNA

Metadata: samples included in analysis.

WGCNA: Gene‑level output from the WGCNA including Ensembl gene ID, mouse gene symbol, gene description, and assigned module color. For each gene, normalized and batch‑corrected expression values (log2 CPM) are provided across all samples. Module color assignments reflect the final merged module structure used for downstream module–trait correlations, hub gene identification, and functional enrichment analyses. Genes assigned to the “grey” module were not included in network modules.

Genes per Module: number of genes in each module

Correlations: Pearson’s correlations (R and FDR corrected pvalues) for correlations between each ME and Diet, LPS, LPS x Diet and sex.

GO Terms: Significantly enriched GO terms within each module for Biological Process (BP), Cellular Component (CC) and Molecular Function (MF).

MGEnrichment: Output of significantly enriched lists from MGEnrichment database for blue and black module.

S3. DEGs. **Differentially expressed genes (DEGs) across experimental conditions.**

**RawCounts: raw, un-normalized counts for each gene (rows) and sample (columns)**

**Metadata: Animal and group information for each sample.**

**Statistics:** gene‑level differential expression results for each comparison. Log Fold Change (LogFC), t values, p pvalues (P.Value) and FDR adjusted pvalues (adj.P.Val), beta values (B), direction of expression change relative to vehicle controls and Significance (NonDE vs DE) for each comparison.

DEG counts: Number of differentially expressed genes for each comparison.

Per Sample Values: RPKM and Log2CPM values for each gene and sample.

S4. GSEA. **Gene set enrichment analysis (GSEA) summary of microglial transcriptomic signatures across experimental conditions.** GSEA results assessing the enrichment of published microglial gene sets from MGEnrichment across multiple experimental comparisons, including inflammatory stimulation (e.g., LPS vs PBS), short‑chain fatty acid treatments (acetate, butyrate, propionate), sex, genotype, developmental stage, disease models (e.g., Alzheimer’s disease, ALS), injury, and microbiome status. For each comparison, the table reports the gene set (pathway or signature), source study, experimental context, tissue and species, normalized enrichment score (NES), enrichment score (ES), log fold change (logFC), nominal p value, false discovery rate–adjusted p value (FDR), gene set size, and leading‑edge genes driving enrichment. Positive NES values indicate enrichment in the first condition listed, whereas negative NES values indicate depletion.

S5. Morphology. Microglial Morphology Results.

AOVA within Region. ANOVA results for Cluster * Diet *Sex main effects and interactions within each Brain Region. Chi Squared values, degrees of freedom (Df) and P values listed.

Posthocs Diet within LPS: Sidak corrected posthoc comparisons comparing Diet conditions within LPS treatments for each cluster and sex.

Posthocs LPS within Diet: Sidak corrected posthoc comparisons comparing LPS conditions within Diet treatments for each cluster and sex.

Microglial Density Results. ANOVA. ANOVA results Cluster * Diet *Sex main effects and interactions within each Brain Region. Chi Squared values, degrees of freedom (Df) and P values listed.
Density Posthocs: Sidak corrected posthoc comparisons comparing Diet conditions within LPS treatments and Sex.

S6. In vitro assay results.

ANOVA + Tukey Post Hoc results from Multiplex assay for (i) IL10, (ii) IL6, (iii) TNFa and (iv) Il1b. ANOVA + Tukey Post Hoc Results from Griess Reagent Assay, BrdU proliferation assay, Phagocytosis assay and XTT assay.

S7. HDACi and qFLOW results.

ANOVA in vitro qFLOW. *Diet *LPS and *Interaction main effects. F value and P value. qFLOW Sidak Posthocs In vitro: Sidak corrected posthoc comparisons comparing Diet conditions within LPS treatments.

ANOVA in vivo qFLOW. Main effect of *Interaction. F value and P value presented. qFLOW Sidak Posthocs In vivo: Sidak corrected posthoc comparisons comparing Diet conditions within LPS treatments and Sex.

Non Linear Regression of HDAC-Glo Assay Statistics on Log2 of SCFA Concentration. IC50 presented in M.
